# Novelty gates memory formation through ventro-dorsal hippocampal interaction

**DOI:** 10.1101/673871

**Authors:** Felipe Fredes, Maria Alejandra Silva, Peter Koppensteiner, Kenta Kobayashi, Maximilian Joesch, Ryuichi Shigemoto

**Author notes:** Correspondence to: Felipe Fredes or Ryuichi Shigemoto. These authors share senior authorship (Maximilian Joesch, Ryuichi Shigemoto).

## Abstract

Novelty facilitates formation of memories. The detection of novelty and storage of contextual memories are both mediated by the hippocampus, yet the mechanisms that link these two functions remain to be defined. Dentate granule cells (GCs) of the dorsal hippocampus fire upon novelty exposure forming engrams of contextual memory. However, their key excitatory inputs from the entorhinal cortex are not responsive to novelty and are insufficient to make dorsal GCs fire reliably. Here we uncover a powerful glutamatergic pathway to dorsal GCs from ventral hippocampal mossy cells (MCs) that relays novelty, and is necessary and sufficient for driving dorsal GCs activation. Furthermore, manipulation of ventral MCs activity bidirectionally regulates novelty-induced contextual memory acquisition. Our results show that ventral MCs activity controls memory formation through an intra-hippocampal interaction mechanism gated by novelty.

## Introduction

Novelty induces psychological stress and arousal(Feenstra et al., 1995), which often results in efficient memory formation in the daily life(Fenker et al., 2008). Such emotional signaling involves the hippocampus (Kaplan et al., 2014; Knight, 1996) and especially its ventral moiety (Floriou-Servou et al., 2018), whereas encoding of spatial memory has been extensively studied in the dorsal hippocampus (Buzsáki and Moser, 2013). However, functional distinction and interaction between the dorsal and ventral hippocampus largely remain elusive. Granule cells (GCs) in the dorsal dentate gyrus (DG) increase their activity during novel environment exploration(Sariñana et al., 2014) forming engrams of contextual memory(Bernier et al., 2017; Liu et al., 2012; Ramirez et al., 2013; Redondo et al., 2014). They receive major excitatory inputs from the entorhinal cortex in the outer two thirds of molecular layer (ML), whereas another excitatory synaptic connection is made in the inner one third of ML by mossy cells (MCs) located in the hilus (Figure 1A). However, both of these excitatory inputs also innervate interneurons in the hilus(Scharfman, 2016), and rather exert net inhibitory effects on GCs through robust feedforward inhibition(Hsu et al., 2016) (Figure 1A). Furthermore, entorhinal cortex neurons are not responsive to environmental novelty(Burgalossi et al., 2014). Thus, the mechanism underlying initiation of GCs firing that leads to memory engrams formation is still unclear. MCs in the ventral hippocampus have extensive bilateral projections to the dorsal DG(Blasco-Ibáñez and Freund, 1997; Fujise et al., 1997) (Figure 1A). Based on this anatomical finding and the preferential signaling of novelty in the ventral hippocampus(Floriou-Servou et al., 2018), we hypothesized that this long range excitatory pathway could convey novelty signals providing a critical excitatory input to dorsal GCs(Scharfman, 2016), facilitating contextual memory formation.

**Figure. 1.**
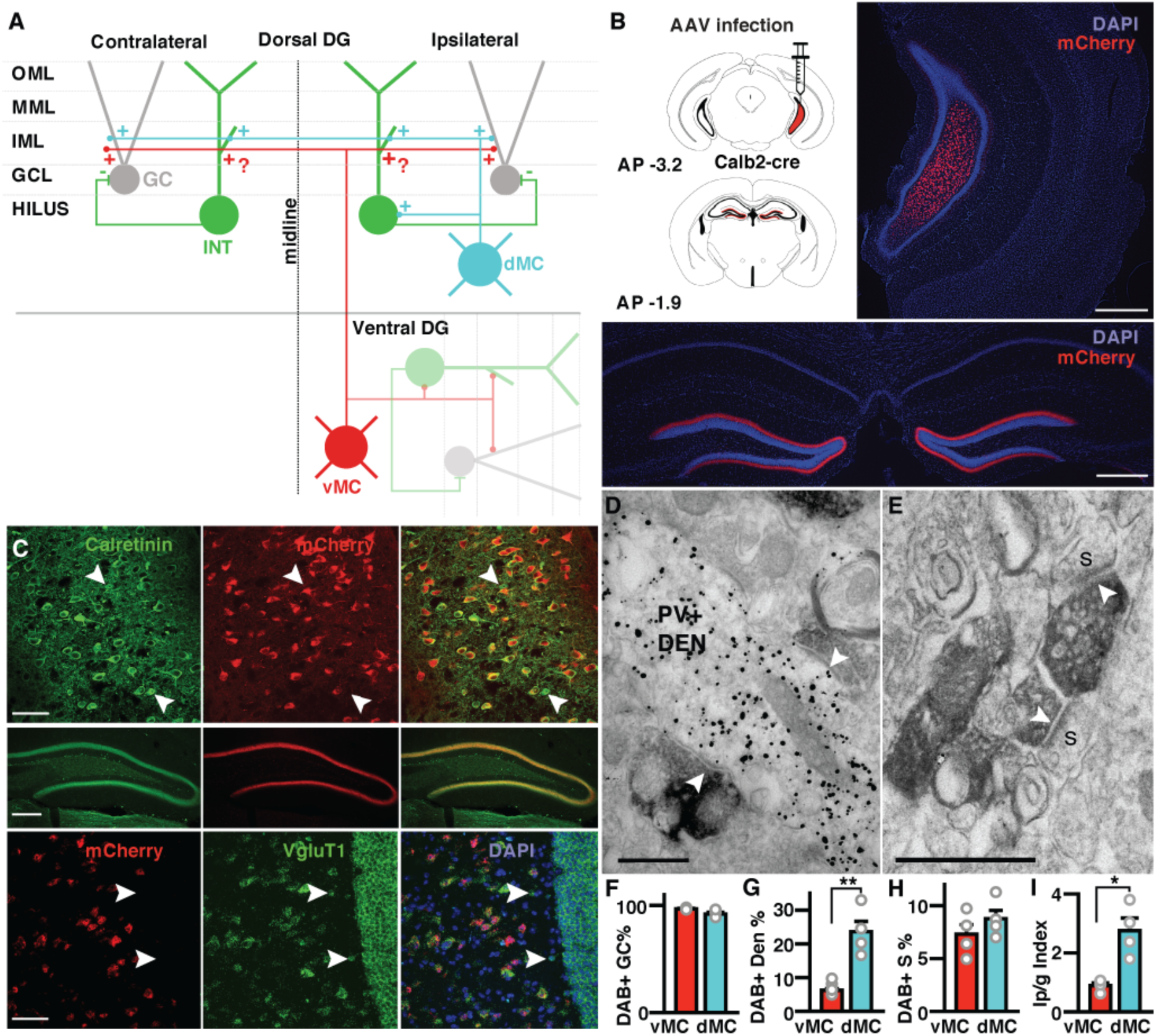
Ventral mossy cells project bilaterally to the dorsal dentate gyrus and mostly contact dentate granule cell spines. **A**. Schematic representation of connectivity between dorsal MCs (cyan) and ventral MCs (red) with interneurons (green) and GCs (grey). IML, inner molecular layer. MML, medial molecular layer. OML, outer molecular layer. GCL, granule cell layer. **B**. Cre-dependent mCherry expression in the ventral hilus and dorsal dentate gyrus after injection of AAV5-hSyn-DIO-mCherry into the ventral hilus of a Calb2-cre mouse. AP, antero-posterior coordinates with respect to bregma. DAPI (4’,6’-diamino-2-phenylindole) labels cell nuclei. Scale bars: 500 µm **C**. mCherry-expressing cells in the ventral hilus and their terminals in the dorsal DG are Calb2 immuno-positive. White arrowheads show Calb2-positive cells that do not express mCherry. Scale bar upper panel: 100 µm, middle panel: 200 µm. Lower panel shows a fluorescent in-situ hybridization for mCherry and vGluT1. Scale bar: 50 µm. White arrows indicate vGluT1-positive cells negative for mCherry. **D**. Two diaminobenzidine (DAB)-labeled terminals from dorsal MCs making synaptic contacts (white arrowheads) with a single PV-positive dendritic shaft (PV+ DEN) immunolabeled with gold particles. **E**. DAB-labeled terminals from ventral MCs synapsing (white arrowheads) onto GCs spines (s). Scale bars: 500 nm. **F**. Quantification of DAB-labeled synapses made on GCs spines (in % of all DAB-labeled synapses) by dorsal MCs (dMC) and ventral MCs (vMC). Mean + s.e., n = 4 animals. **G**. Quantification of dorsal and ventral MCs synapses onto parvalbumin (PV)-positive interneurons. Number of DAB-labeled synapses (in % of total synapses) onto PV-positive dendrites (PV+ DEN). Mean + s.e., n = 4 animals, ** p = 0.003 (Unpaired, two tailed Student’s t-test). **H**. Number of DAB-labeled synapses (in % of total synapses) onto GCs spines (PV+ S) coming from either ventral or dorsal MCs. Mean + s.e., n = 4 animals. **I**. Index Ip/g (see Method) for ventral MCs and dorsal MCs, showing stronger innervation of PV-positive dendrites by dorsal than ventral MCs. Mean + s.e., n = 4 animals, p = 0.006 (Unpaired, two tailed Student’s t-test).

Following this hypothesis, we show in this study that in contrast to dorsal MC projections into the inner one-third of the ML, ventral MC projections provide a powerful net excitatory effect on dorsal GCs. Inhibiting this ventral MC pathway, but not dorsal MC pathway, during the learning phase of fear conditioning in a novel environment abolishes the formation of contextual memory. Furthermore, while animals are unable to form contextual memories when conditioned in a familiar environment, chemogenetic reactivation of ventral MCs is sufficient to rescue the formation of contextual memories in a familiar environment, indicating that information about novelty is relayed to dorsal GCs via ventral MCs. Finally, we show that the population activity of both, ventral MCs and dorsal GCs, is directly linked with the level of environmental novelty during exploration - it decreases with familiarization and rebounds when animals explore a new environment again. Thus, our data unveil a mechanistic link between novelty detection and memory formation by direct interaction of the ventral and dorsal hippocampus.

## Results

### Ventral MCs make extensive excitatory synaptic contacts with dorsal GCs

MCs in the ventral, but not dorsal region of the hippocampus are the only calretinin (Calb2)-positive neurons in the DG(Blasco-Ibáñez and Freund, 1997; Cembrowski et al., 2016; Fujise et al., 1997) enabling targeted manipulations of ventral MCs using Calb2-cre mice. After injection of cre-dependent adeno-associated virus (AAV) expressing mCherry into the ventral hilus of a Calb2-cre mouse (Figure 1B), we identified a dense and bilateral projection to the inner ML of the dorsal DG that extensively spanned the whole dorso-ventral hippocampal axis (Figure 1B) with no other distinct projections in the hippocampus (Figure S1A). The vast majority of mCherry-expressing cells in the ventral hilus (92%, n = 175 cells), as well as their terminals in the dorsal DG, were Calb2-positive as expected (Figure 1C). Virtually no cells in the granule cell layer and only a small number of pyramidal cells in the CA3 area (7.3% of all labeled cells) expressed mCherry (Figure S1A, B). *In situ* hybridization showed that all the mCherry-expressing cells are vGluT1-positive (n = 67 cells, Figure 1C) confirming that they are glutamatergic. Given that both dorsal and ventral MCs project to the dorsal inner ML (Figure 1A) and that optogenetic stimulation of dorsal MCs axons primarily drives interneuron firing(Hsu et al., 2016), we compared the postsynaptic targets of ventral and dorsal MCs in the dorsal inner ML. The majority of interneuron dendrites in ML originate from parvalbumin-positive (PV+) basket cells (85 %, Figure S2H-J), which are robustly activated by dorsal MCs(Hsu et al., 2016), have the highest probability of innervating GCs among all interneurons and thus have a strong direct influence on GC activity(Espinoza et al., 2018). Therefore, we quantified by electron microscopy the numbers of asymmetrical (glutamatergic) synapses arising from ventral or dorsal MCs onto dendritic shafts of PV+ interneurons and those onto GCs spines within the inner ML of dorsal DG (Figure 1D-I, also see Methods). To visualize MCs terminals, we expressed a membrane-bound fluorescent protein in ventral and dorsal MCs and immunolabeled their terminals with diaminobenzidine (DAB) (Figure 1D, E). Asymmetrical synapses made by dorsal MCs terminals were identified in the inner ML by AAV injection into the contralateral hilus(Hsu et al., 2016) (Figure S2A-C). Then, we immunolabeled PV+ interneurons with gold particles (Figure 1D, also see Methods and Figure S2A-G)). The vast majority of DAB-positive terminals (>97 % for ventral MCs, >92 % for dorsal MCs, Figure 1F. Also see Methods and Table S1) made synapses with GC spines (Figure 1E), whereas small numbers of DAB-positive terminals with PV+ dendritic shafts (Figure 1D). No PV+ spines (n = 20) made synapses with DAB-positive terminals (Figure S2F-G). On the PV+ dendritic shafts, we found a higher percentage of DAB-positive synapses made by dorsal (23.6 %) than ventral (6.22 %) MCs (Figure 1G, p = 0.003), whereas no significant difference was observed on GCs spines (Figure 1H). To control for any bias in MC infection rates, we also compare the relative innervation of PV+ interneurons normalized by that of GCs (Ip/g index, see Methods), and found a 3.1-fold higher Ip/g value for dorsal than ventral MCs (Figure 1I, p = 0.006). Given that only a partial ipsilateral population of dorsal MCs is labeled in our experiments, the high percentage (23.6 %) of labeled synapses on PV+ dendrites implies that the majority of excitatory inputs on these interneurons in the inner ML derive from dorsal MCs, and only a minor population from ventral MCs. Thus, ventral MCs may exert a net excitatory effect on dorsal GCs in contrast to the net inhibitory effect exerted by dorsal MCs(Bui et al., 2018; Hsu et al., 2016).

### Ventral MCs activity is required for dorsal GCs activation during novel environment exploration

To test if the ventral MCs are capable to influence novelty induced changes in the dorsal hippocampus, we modulated ventral MCs activity during novel environment exploration and assayed changes in immediate early gene (c-Fos) expression in GCs, a proxy for neuronal activity. We injected AAV expressing inhibitory designer receptors exclusively activated by designer drugs (DREADDs) (Hm4Di) bilaterally into the ventral hilus of Calb2-cre mice. We confirmed that the vast majority of Hm4Di-expressing cells were Calb2-positive (95 %, n = 134 cells). Three weeks later these animals were injected with clozapine-N-oxide (CNO) intraperitoneally (i.p.) 40 min before novel environment exploration (Figure 2A). We found a significantly reduced number of c-Fos expressing dorsal GCs in the upper blade in animals expressing Hm4Di compared to the control group expressing only mCherry (Figure 2B, C, G). The lower blade showed no significant reduction in c-Fos expression (Figure 2G). In contrast, using the same procedure in animals expressing excitatory DREADDs (Hm3Dq), we found a 6-fold increase in c-Fos-positive GCs in the upper blade of the dorsal DG (Figure 2B, D, H). Within the ventral DG of these animals, we also observed an increase of c-Fos-positive GCs compared to mCherry-expressing animals, but the increment was much less robust compared to that in the dorsal DG (Figure 2E, F, H). We also found an increased c-Fos expression in ventral hilar somatostatin-positive interneurons in Hm3Dq-expressing animals compared to mCherry-expressing controls (42 % in Hm3Dq n = 96 cells, versus 0 % in mCherry) (Figure 2E, F). To examine if the effects of inhibitory and excitatory DREADDs are specific to ventral but not dorsal MCs, we injected AAVs expressing mCherry, Hm4Di-mCherry or Hm3Dq-mCherry bilaterally into the dorsal hilus of Drd2-cre mice expressing Cre in dorsal MCs(Puighermanal et al., 2015). We confirmed that all mCherry-expressing cells are positive for GluR2/3, a marker for dorsal MCs(Fujise et al., 1997) (Figure S5A-C, n = 85 cells). Novel environment exploration after CNO injection showed significant reduction of c-Fos-positive dorsal GCs in Hm3Dq-but not Hm4Di-expressing animals (Figure S3), consistent with the net inhibitory effect on dorsal GCs exerted by dorsal MCs. The excitatory effects of Hm3Dq in ventral (Figure S4A-E) and dorsal MCs (Figure S4F-J) were confirmed by measuring firing and membrane potentials. In contrast with the inhibitory effects of dorsal MCs activation on dorsal GCs(Bui et al., 2018; Hsu et al., 2016), these results indicate that the activation of long-range projections of ventral MCs has an excitatory net effect on dorsal GCs and further support our anatomical findings.

**Figure. 2.**
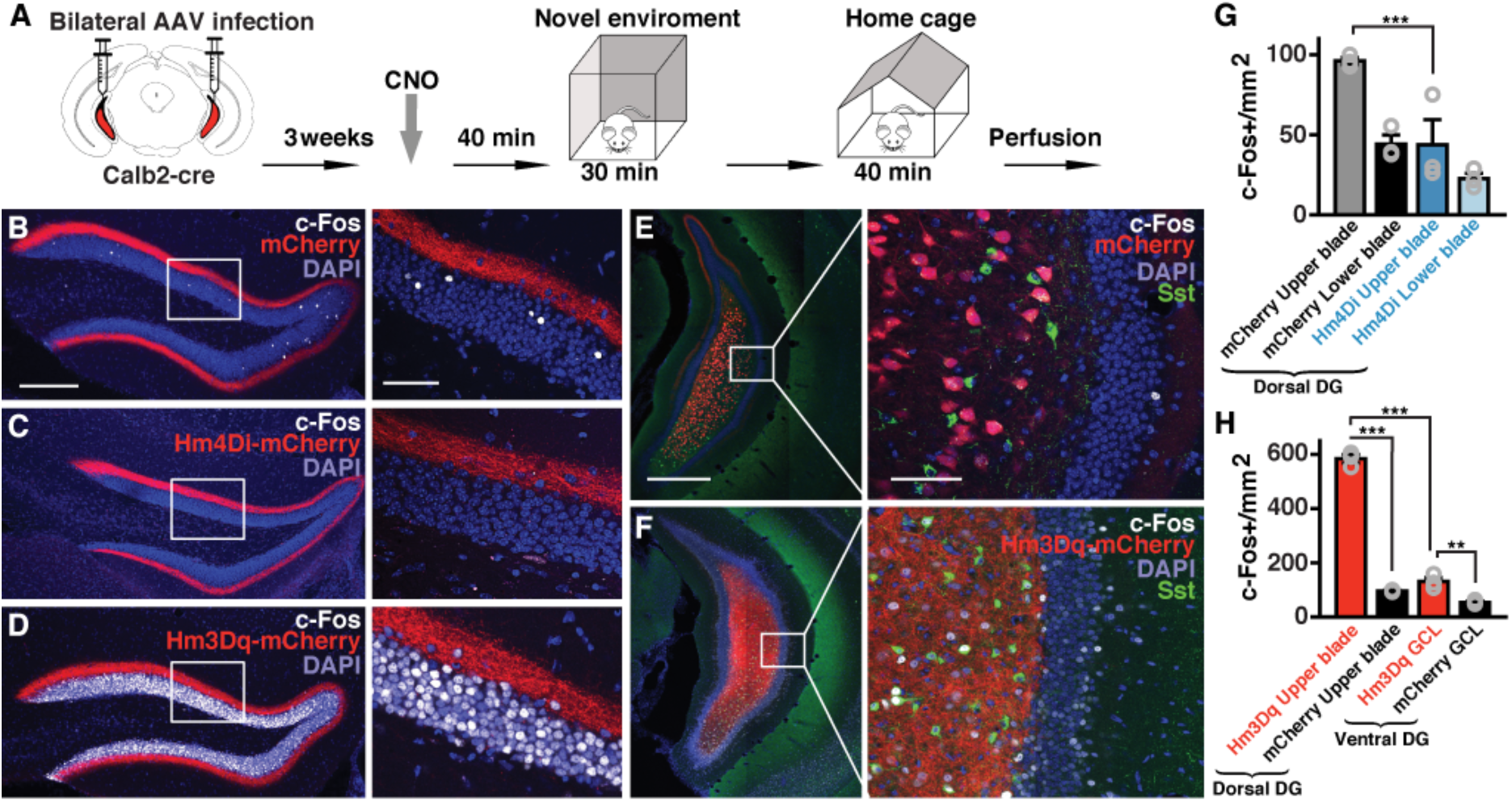
Ventral mossy cell activity is necessary and sufficient to activate dorsal granule cells. **A**. Schematic of experimental procedures where Calb2-cre mice were injected bilaterally with AAVs expressing mCherry, inhibitory DREADDs (Hm4Di-mCherry) or excitatory DREADDs (Hm3Dq-mCherry) in the ventral hilus. **B-D**. Confocal images of the dorsal DG showing c-Fos immunoreactivity (white) from animals expressing mCherry (**B**), Hm4Di-mCherry (**C**) or Hm3Dq-mCherry (**D**). Higher magnification images of the boxed areas (left) are shown on the right. Scale bars: 200 µm (left), 100 µm (right). **E-F.** Confocal images of the ventral DG showing c-Fos immunoreactivity from animals expressing mCherry (**E**) or Hm3Dq-mCherry (**F**). Scales bars: 500 µm. Insets show a higher magnification of the boxed areas. Scale bar: 100 µm. Somatostatin (Sst) immunopositive cells and terminals are shown in green. **G**. Quantification of c-Fos-positive GCs in the upper and lower blades of the dorsal DG from animals expressing mCherry or Hm4Di-mCherry (n = 3 animals per group; mean + s.e. *** p < 0.001, t-test). **H**. Quantification of c-Fos-positive GCs in the upper blade of the dorsal DG and GC layer (GCL) in the ventral DG from animals expressing mCherry or Hm3Dq-mCherry (n=3 animals per group; mean + s.e. *** p < 0.001, ** p = 0.008, t-test).

### Ventral MCs are a powerful drive of GC activity *in vivo*

To further examine the strength of the excitatory effect of ventral MC projections over the dorsal GCs *in vivo*, we expressed Hm3Dq bilaterally in the ventral MCs and the calcium indicator GCaMP6s unilaterally in the dorsal GCs in double transgenic mice Prox1-creERT2/Calb2-cre (Figure 3A). Through a GRIN lens implanted over the dorsal DG, we found the activity of dorsal GCs dramatically increased 30 min after CNO injection during familiar environment exploration (Figure 3B, C. See also Movie S1). This drastic change in activity has no apparent correlation with locomotion speed, which are comparable in Hm3Dq-expressing and control animals (Figure 3D, E). The overall strength of the measured responses indicate that ventral MCs activity can have a critical role in driving dorsal GCs firing, suggesting a directed flow of information from the ventral to the dorsal hippocampus.

**Figure 3.**
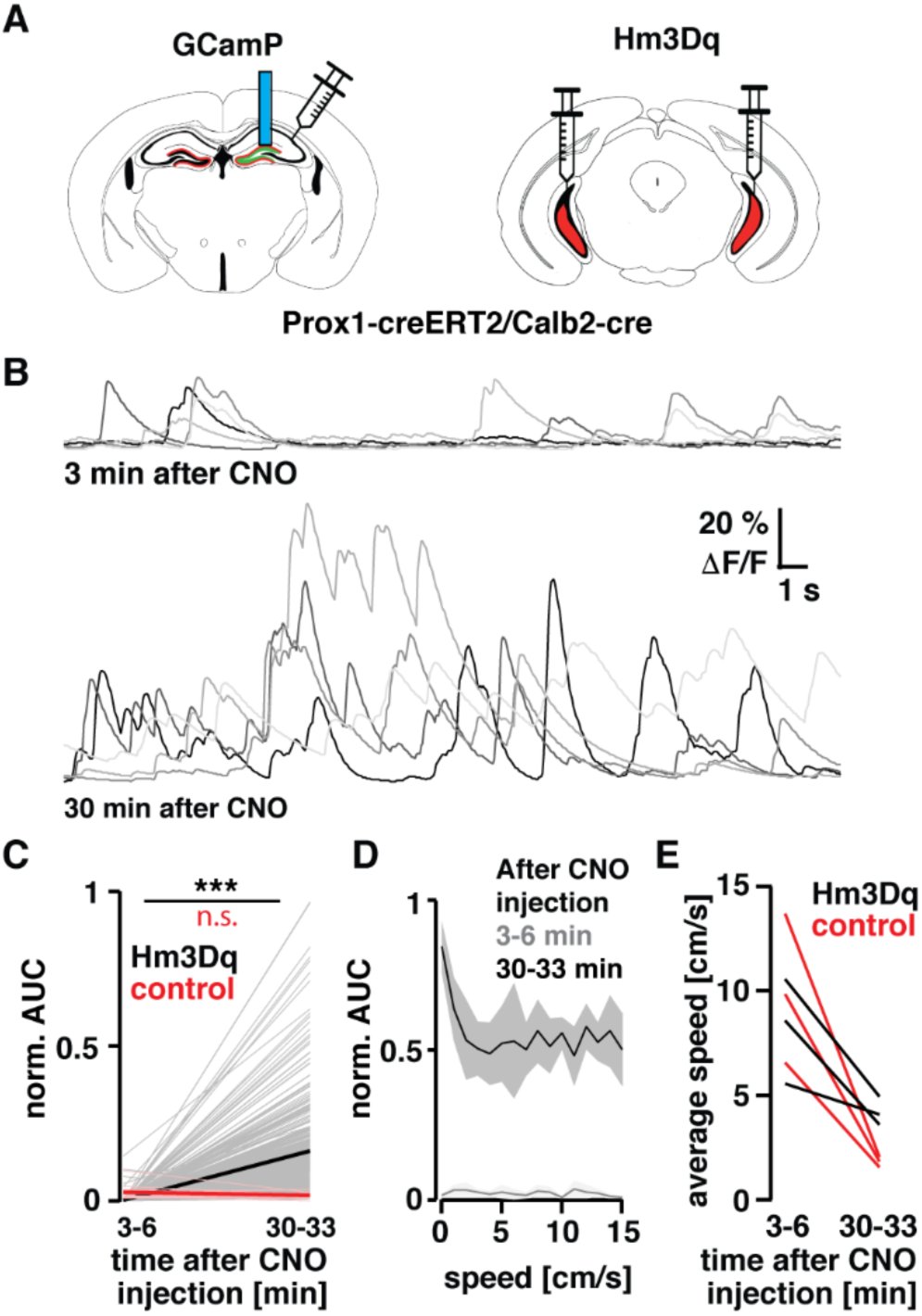
Ventral MCs activation exerts a strong influence on dorsal GC activity *in vivo*. **A**. Schematic of experimental procedures where double transgenic mice Prox1-creERT2/Calb2-cre were injected bilaterally with AAV expressing Hm3Dq-mCherry in the ventral hilus, and unilaterally with AAV expressing GCaMP6s in the dorsal DG. Later a GRIN lens (blue rod) was implanted above the injection site in the dorsal DG. **B**. Examples of dorsal GC calcium transients before and after the effect of CNO. Note: the five most active cells are presented on the top traces, whereas a random set is selected in the bottom. **C**. Change in mean activity (AUC – see methods) for all recorded cells before and after the effect of CNO, 3-6 and 30-33 min after CNO injection, respectively (Hm3Dq: n = 386 cells, p < 0.001 Wilcoxon signed-rank test, Control n = 15 cells, not significant). Note: the reduced number of cells in the control condition reflects the normal sparse firing of dorsal GCs. **D**. Average activity distribution in Hm3Dq-expressing animals before and after the effect of CNO (as in **C**) binned by locomotion. **E**. Average locomotion speed of Hm3Dq-expressing and control animals. Note that the higher initial speed appears to be a distress response to the injection.

### Ventral MCs activity is required for context memory formation

Considering that dorsal GCs activity is necessary for contextual memory formation(Kheirbek et al., 2013; Ramirez et al., 2013; Redondo et al., 2014), we examined if ventral MCs activity is required for context-dependent memory formation in fear conditioning. We expressed Hm4Di bilaterally in the ventral MCs of Calb2-cre mice (Figure 4A) and injected CNO i.p. 40 mins before fear conditioning (Figure 4B). These animals showed significantly reduced freezing levels (Figure 4E) compared to the mCherry expressing control group, indicating that ventral MCs activity is necessary for context memory formation. Since acute ablation of dorsal MCs was reported to impair context discrimination(Jinde et al., 2012), we next tested the effect of inhibitory DREADDs in dorsal MCs using Drd2-cre mice (Figure 4A). These animals showed significantly increased freezing levels in retrieval (Figure 4E) indicating distinct roles of ventral and dorsal MCs. Although these animals also tended to increase freezing in conditioning (Figure 4D), which may affect memory acquisition, the increased freezing in retrieval was context-dependent (Figure 4E-F). None of the experimental groups showed differences compared to control animals during preexposure or different context (Figure 4C, F). Using acute slice preparation from Drd2-cre animals expressing channelrhodopsin (ChR2) and Hm4Di in dorsal MCs, we confirmed that light-evoked monosynaptic excitatory and disynaptic inhibitory responses recorded in dorsal GCs(Hsu et al., 2016) was significantly reduced by CNO (3 μM) (Figure S5D-F). These results indicate that ventral MC activity, but not dorsal MC activity, is indispensable for context memory formation.

**Figure 4.**
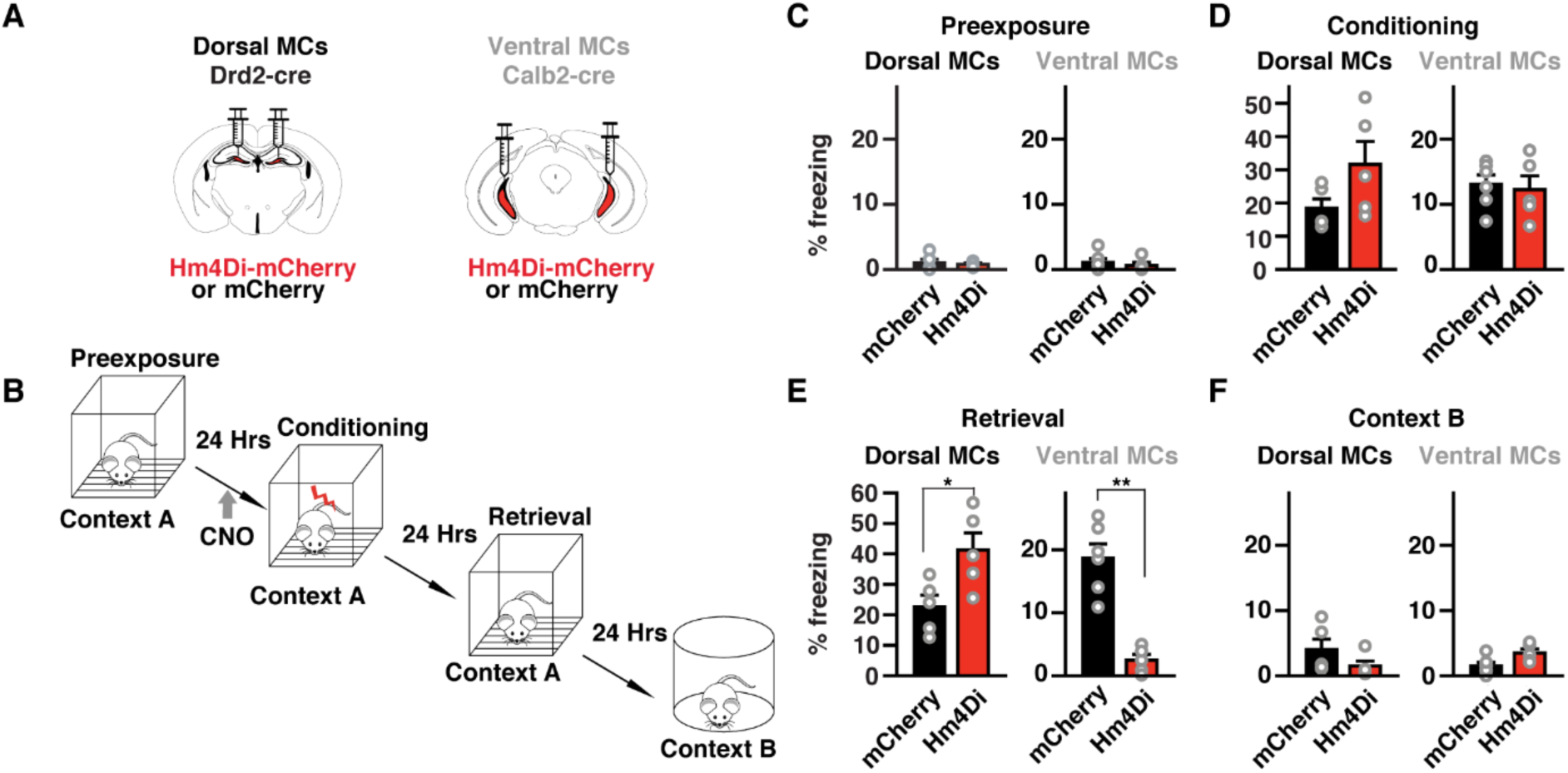
Differential role of dorsal and ventral MCs in context memory formation. **A**. Schematic representation of the injection procedure in Drd2-cre and Calb2-cre mice. **B**. Schematic representation of the behavioral paradigm. **C-F**. Freezing levels during pre-exposure (**C**), conditioning (**D**), retrieval (**E**) and exposure to a novel context B (**F**). Scale bar: 200 µm. * p = 0.032. ** p = 0.004 Mann-Whitney U-test. For dorsal MCs experiments n = 5 mice for Hm4Di and n = 5 mice for mCherry. For ventral MCs experiments n = 5 mice for mCherry and Hm4Di. n = 6 mice for Hm3dq.

### Reactivation of ventral MCs rescues context memory formation

Familiarization is known to reduce arousal(Feenstra et al., 1995). Thus, we hypothesized that animals in a familiar environment should form contextual memory less efficiently than in a novel one(Hall et al., 2000). To test this, we familiarized animals in the conditioning cage for 10 mins, during 6 consecutive days and then performed fear conditioning in the same cage on day 6. As expected, these animals showed significantly lower freezing levels in the retrieval session than animals conditioned in a novel environment (Figure 5D). To examine if ventral MCs activation can induce contextual memory formation in a familiar environment, we bilaterally expressed Hm3Dq in ventral MCs and titrate the amount of CNO necessary to cause only a slight increase of dorsal GCs activity. We found that as little as 0.75 mg/kg of CNO (i.p.) roughly doubled the number of c-Fos-positive dorsal GCs when animals explored a novel environment (Figure 5B, C). Then, we bilaterally expressed Hm3Dq in ventral MCs and familiarized these animals in the conditioning chamber as described before and injected 0.75 mg/kg of CNO 30 min before conditioning (Figure 5A). We found that freezing levels during retrieval were rescued to comparable levels of animals conditioned in a novel environment (Figure 5D). This rescue effect was context dependent since no increase in freezing was observed in context B (Figure 5D). These results indicate that ventral MCs activity is sufficient to form context memory in a familiar environment.

**Figure. 5.**
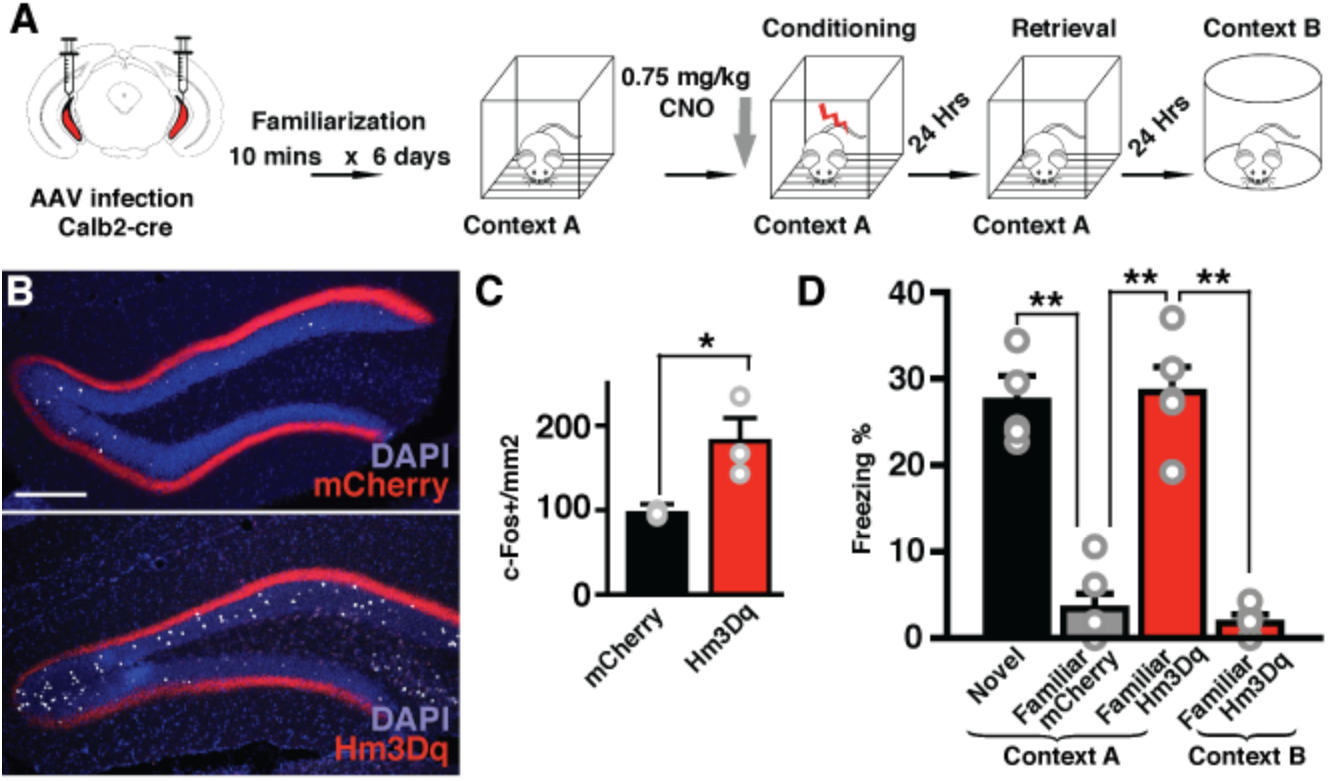
Activation of ventral MCs recovers novelty induced memory formation. **A.** Schematic representation of the familiarization fear conditioning paradigm and ventral MC manipulation. **B**. CNO titration for memory enhancement experiments using animals injected with 0.75mg/kg of CNO. Representative confocal images of the dorsal DG of animals expressing mCherry or Hm3Dq-mCherry in ventral MCs, showing c-Fos-positive dorsal GCs in white. Scale bar, 200 µm. **C**. Quantification of c-Fos-positive dorsal GCs (* p = 0.036, t-test. n = 3 animals for mCherry (same data from Figure 2B, G), n = 3 for Hm3Dq-mCherry). **D.** Effect of ventral MCs excitation by 0.75 mg/kg of CNO on freezing levels during the retrieval. Fear conditioning in a familiar environment did not cause freezing during the retrieval session (familiar), but excitation of ventral MCs restored freezing (familiar Hm3Dq) to a similar level as obtained with conditioning in a novel environment (novel). This manipulation did not have any effect in freezing levels in a new environment (Context B). (** p < 0.01, Mann-Whitney U-test. n = 6 animals for novel, n = 5 animals for familiar and familiar Hm3Dq).

### Long-range axonal projections of ventral MCs onto dorsal MCs are necessary for contextual memory formation

Ventral MCs project bilaterally via long-range projection to dorsal GCs, but also have local axonal arborization within the ventral hippocampus. To conclusively determine if the long-range projections are specifically required for context memory formation, we perturbed synaptic transmission of ventral MC projections while animals were conditioned in a context learning paradigm (Figure 6). For this purpose, we took advantage that Hm4Di mainly blocks synaptic release at axon terminals(Stachniak et al., 2014) with no effect on firing rate at cell bodies, and local intracerebral injections of CNO can selectively inhibit one pathway without affecting the other collaterals(Stachniak et al., 2014). Using acute slice preparation from Calb2-cre animals expressing channelrhodopsin (ChR2) and Hm4Di in ventral MCs, we confirmed that CNO (3 μM) significantly reduces light-evoked EPSC amplitudes in dorsal GCs but not firing of ventral MCs (Figure S6). To functionally dissect the role of axonal projections from ventral MCs during fear conditioning, we bilaterally expressed Hm4Di or mCherry in ventral MCs, and implanted a bilateral cannula over the dorsal DG (Figure 6B). Animals then underwent fear conditioning (Figure 6A) and received a small volume of CNO (3 μM, 300 nl) delivered directly into the dorsal DG bilaterally through the cannulas. If we administered CNO 15 min before conditioning, animals expressing Hm4Di showed a robust reduction in freezing during retrieval compared to mCherry expressing control animals (Figure 6E). However, infusing CNO right after conditioning or before the retrieval had no effect on freezing compared to controls (Figure 6E). We then tested if inhibiting local hilar ventral MCs terminals in the ventral DG altered memory acquisition by delivering CNO locally through cannulas implanted bilaterally into the ventral hilus (Figure 6B). This manipulation had no effect on freezing compared to control animals (Figure 6E). None of these manipulations had any significant effect on freezing levels during pre-exposure (Figure 6C), conditioning (Figure 6D) or exposure to a different environment, context B (Figure 6F). These results indicate that the activity of ventral MC terminals in the inner ML of the dorsal DG drives dorsal GCs firing and is necessary for contextual memory acquisition.

**Figure. 6.**
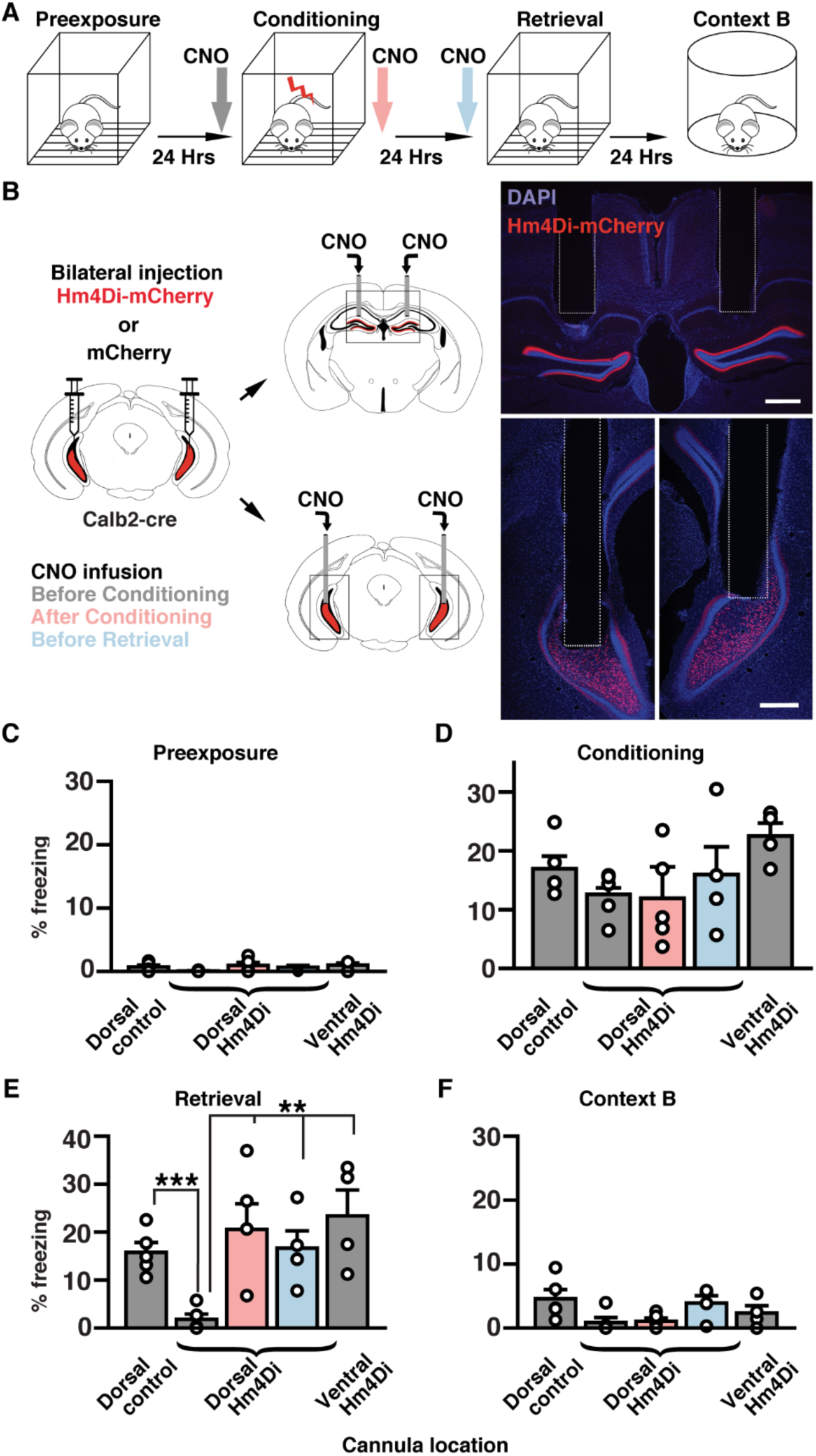
Ventral mossy cell terminal activity in the dorsal DG is indispensable for fear context memory formation. **A**. Fear conditioning protocol. CNO was infused before conditioning (gray), after conditioning (pink), or before retrieval (blue). **B**. Calb2-cre mice were injected bilaterally with AAVs expressing Hm4Di-mCherry or mCherry into the ventral hilus, and CNO was infused into the bilateral dorsal or ventral DG. Photographs (right) show examples of cannula placement. Scale bars: 500 µm. **C**. Percentage of freezing during preexposure shows no statistical differences (p > 0.05, Mann-Whitney U-test) among five groups, dorsal control group expressing mCherry and receiving CNO in the dorsal DG before conditioning (n = 5 animals), dorsal Hm4Di groups expressing Hm4Di-mCherry and receiving CNO in the dorsal DG before conditioning (gray, n = 5 animals), after conditioning (pink, n = 5 animals) or before retrieval (blue, n = 4 animals), and ventral Hm4Di group expressing Hm4Di-mCherry and receiving CNO in the ventral DG before conditioning (n = 4 animals). **D**. Percentage of freezing during conditioning shows no significant differences among all groups (p > 0.05, Mann-Whitney U-test). **E**. Percentage of freezing during retrieval. Dorsal Hm4Di group receiving CNO before conditioning shows a robust reduction in freezing compared to the dorsal control group (*** p < 0.001, Mann-Whitney U-test), but not other dorsal Hm4Di groups receiving CNO after conditioning or before retrieval, and ventral Hm4Di group (** p < 0.01, Mann-Whitney U-test). **F**. Percentage of freezing during exposure to context B shows no significant differences among all groups (p > 0.05, Mann-Whitney U-test).

### Environmental novelty is a strong drive of ventral mossy cells and dorsal granule cells

Our behavioral experiments suggest that ventral MCs relay novelty signals to dorsal GCs, which have been shown to be activated by novel environment exploration(Sariñana et al., 2014). Thus, ventral MCs should also be activated upon novel environment exploration, and activity of ventral MCs and dorsal GCs should diminish in parallel with familiarization of the context. To test this, we recorded neuronal activity of ventral MC and dorsal GC by *in vivo* Ca^2+^ imaging. First, we expressed GCaMP6s in dorsal GCs (Figure S7B) or ventral MCs (Figure S7F) using Prox1-creERT2 or Calb2-cre mice, respectively. We then implanted a GRIN lens coupled to a head-mounted microendoscope to monitor activity-related calcium transients (Figure S7A, E). Animals freely explored a novel environment on day 1 for 20 min and then revisited the same environment for 5 additional days. Ca^2+^ imaging in these freely behaving animals revealed that both, dorsal GCs (Figure 7A-C, Figure S7C, D, M, Movie S2) and ventral MCs (Figure 7F-H, Figure S7G, H, N, Movie S3), showed robust activity during novel environment exploration, and decreased their activity during familiarization (Figure 7C, H, Figure S7M-O). Moreover, the activity of both cell types immediately rebounded upon exposure to a new environment on day 6 (Figure 7C, H, Figure S7D, H, M-N, Movie S2, S3). These changes in activity are driven in both cases mainly by changes in event sizes (Figure S7I-L). Because locomotion increases in novel environment, and could positively correlate with GC and MC activity(Danielson et al., 2017; Pilz et al., 2016), we examined relationship between the speed of mouse movements and activity of dorsal GCs and ventral MCs in novel and familiar environment (Figure 7D-E, I-J, Movie S2, S3). Although the average locomotion speed showed a tendency to decrease through familiarization, this decrease was not significant (Figure 7D, I). Only dorsal GCs activity tended to increase with locomotion speed in the novel (but not in familiar) environment (Figure 7E). However, no apparent increase of ventral MCs activity correlated with the speed was observed (Figure 7J), indicating that the increased activity of ventral MCs in the novel environment is not due to the increased locomotion. To control for other behavioral influences on ventral MC activity, we designed an experiment in which the behavioral statistics remain constant (running on a wheel) across familiarization, and observed twice as much average activity while running in the novel, compared to the familiar environment (Figure 7K). The correlated activity of ventral MCs and dorsal GCs thus suggests that the activity of ventral MCs relay on a population level locomotion-independent novelty signals onto dorsal GCs.

**Figure. 7.**
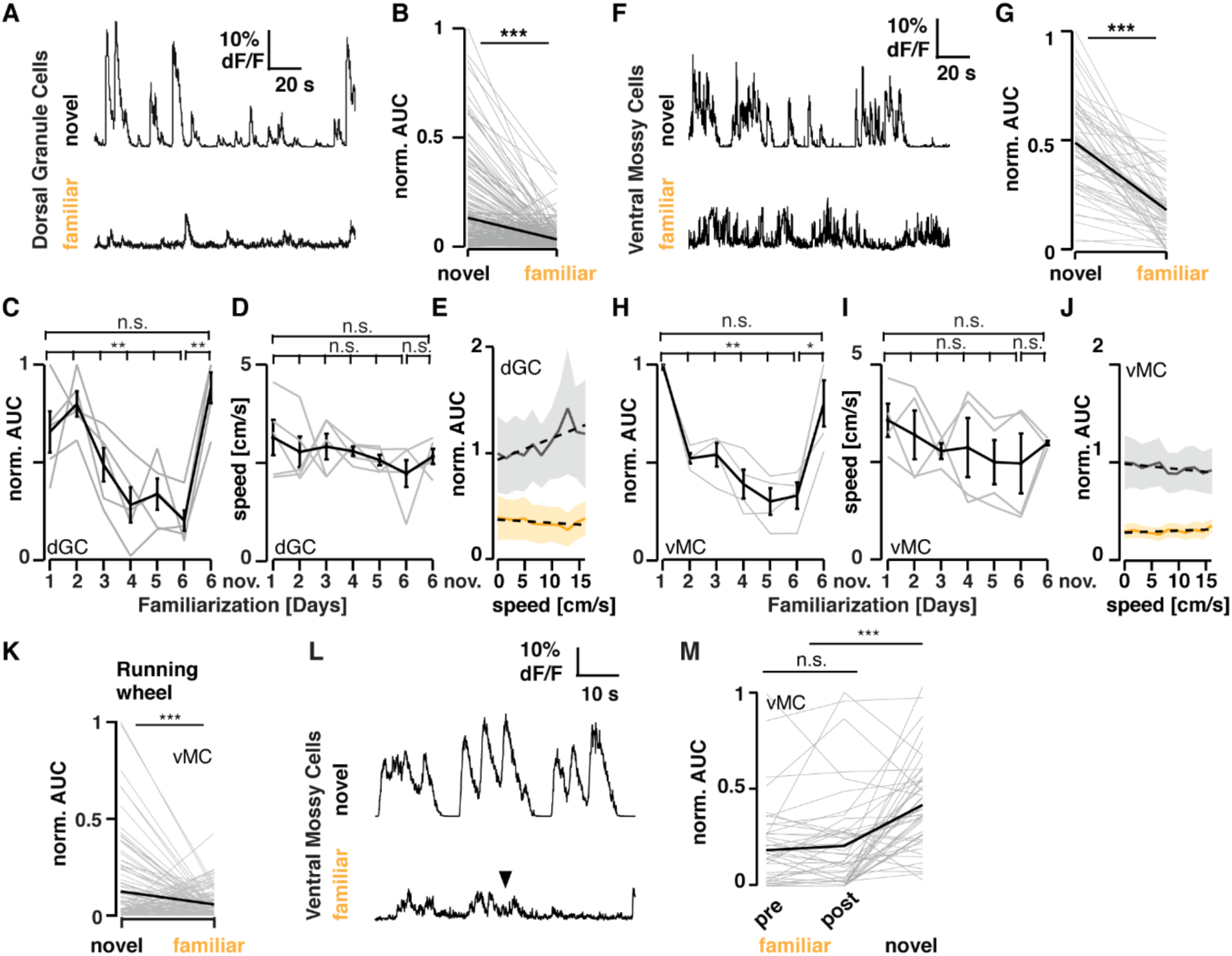
Ventral MCs and dorsal GCs are robustly activated by novel environment exploration. **A.** Example endoscope calcium imaging traces of the same dorsal GC during exploration of a novel and the same familiarized environment 6 days later (Prox1-creERT2 animals unilaterally injected with AAV9-hSyn-Flex-GCaMP6s into the dorsal DG). **B.** Changes in mean activity (AUC – see methods) for all recorded cells from novel (Day1) to the same familiarized (Day 6) environment (n = 229 cells, 5 animals, *** p < 0.001 Wilcoxon signed-rank test). **C.** Normalized average activity for individual (grey trace) and the mean of all animals (black trace +/− SEM) across familiarization and novel environment exploration (5 animals, ** p < 0.01; Kruskal-Wallis one-way ANOVA; Dunn’s multiple comparison test revealed p = 0.08 between Day 1 and Day 6, and p = 0.005 between Day 2 and Day 6). **D.** Average locomotion speed (grey trace) and the mean of all animals (black trace +/− SEM) across familiarization and novel environment exploration (n = 5 animals). **E.** Average neuronal activity relative to locomotion speed in novel (black +/− SEM; day 1 and novel day 6) and familiar (yellow +/− SEM; day 5 and familiar day 6) environments (dotted lines: linear fits, normalized to the maximal response in the first bin). **F-J** as in **A-E**, but with Calb2-cre animals unilaterally injected with AAV5-hSyn-Flex-GCaMP6s and ventral MCs imaged in the ventral hilus. **G**. as in **B** but for ventral MC (n = 60 cells, 4 animals, *** p < 0.001 Wilcoxon signed-rank test). **H**. as in **C** but for ventral MC (4 animals, * p < 0.05; ** p < 0.01, Kruskal-Wallis one-way ANOVA; Dunn’s multiple comparison test revealed p = 0.01 between Day 1 and Day 5, and p = 0.02 between Day 1 and Day 6). **I & J.** as in **D** & **E**, but for ventral MC. **K.** Changes in mean activity (AUC) for all recorded ventral MCs during wheel running epochs in a novel or familiarized environment (n = 110 cells, *** p < 0.001 Wilcoxon signed-rank test). **L**. Representative traces of Ca^2+^ transients of a single cell in familiar and novel environment. Foot shock (inverted triangle) did not cause any changes in Ca^2+^ activity levels in ventral MCs. **M**. Population activity (AUC normalized to the max. AUC of Ca^2+^ transients per animal) of ventral MCs before (pre) and after (post) foot shock in a familiar environment, and that in a novel environment. *** p < 0.001 Mann-Whitney U-test. n = 3 mice.

### Novelty, not surprise, drives MC activity

Novelty is the quality of not being previously experienced. Therefore, the detection of novelty requires examining contents of memory to determine if a stimulus has been experienced in the past, as is the case during environmental exploration. In contrast, the detection of unexpected or surprising events would not (Barto et al., 2013). The foot-shock provided during conditioning, could serve as either a novel or surprising signal, and modulate ventral MC activity. To test this possibility, we performed Ca^2+^ imaging in ventral MCs and familiarized these animals in the conditioning cage, 10 min for 6 days. In accordance with our expectations, ventral MCs were not reactivated by foot-shock but reactivated only with novel environment exploration (Figure 7L, M), indicating that the foot-shock serves as an unexpected or surprising event, rather than novel one in contextual memory formation.

## Discussion

Although hippocampal novelty detection has been shown to be associated with the projections from locus coeruleus (LC) to CA1 and CA3(Takeuchi et al., 2016; Wagatsuma et al., 2017) the mechanisms of novelty-induced activation of the dentate gyrus have not been yet investigated. The increase in activity of dorsal GCs cannot be explained by medial entorhinal inputs, given that they show no change in firing rate with environmental novelty(Burgalossi et al., 2014). In contrast, here we show that ventral MCs have a strong activity increase during novel environment exploration. We propose that this population activity, due to their widespread bilateral projections, can modulate the input relayed from the entorhinal cortex to dorsal GCs by shifting the baseline for depolarization globally. By this mechanism, novelty gates encoding of spatial information relayed by the entorhinal cortex, forming the engram of contextual memory in dorsal GCs. While dorsal MCs have been shown to be involved in spatial coding, being required for the discrimination of changes in local contexts in similar environments(Bui et al., 2018; Danielson et al., 2017; Senzai and Buzsáki, 2017), our results emphasize that their ventral counterparts can relay a global environmental novelty context. Such novelty also activates tyrosine-hydroxylase-expressing LC neurons, which enhances memory consolidation(Takeuchi et al., 2016). However, the LC and ventral MC pathways are apparently independent, because chemogenetic activation of ventral MCs is sufficient to facilitate the formation of contextual memories in familiar environments and contrary to the post-encoding effect of LC activation(Takeuchi et al., 2016), inhibition of ventral MCs right after the conditioning has no effect on memory formation. Although novelty-induced emotional signals could be also involved in the ventral MC activation, it is striking that even electrical shock caused no activation in ventral MCs. Since novelty, by one mean or another, requires the examination of memory contents (Barto et al., 2013), our study opens intriguing new questions regarding the network computations that drive ventral MCs.

Other studies have also tested the role of ventral and dorsal MCs *in vivo* from different perspectives, employing different manipulations and spatiotemporal specificity. A recent study shows that optogenetic stimulation or inhibition of ventral MCs was able to reduce or increase the number of spontaneous epileptic seizures, respectively(Bui et al., 2018). These results are in apparent contradiction with our data as this may imply a net inhibitory effect of ventral MCs on dorsal GCs. However, in contrast to our study, their ventral placement is directed within the intermediate DG (Fanselow and Dong, 2010; Strange et al., 2014). In this area there is a mixture of calretinin-positive and negative MCs that may result in a predominantly inhibitory net effect on the DG after optical stimulation of MCs. In all our experiments we aimed at the ventral one-third of the hilus, where the highest density of calretinin-positive cells is located (Cembrowski et al., 2016). In another study, all MCs were permanently ablated by diphtheria toxin (DT) (Jinde et al., 2012) with two phases, an acute (4-11 days) and a chronic one (4-6 weeks after DT ablation). Dorsal MCs were almost entirely ablated after one week, whereas ventral MCs died only four weeks after DT treatment. Interestingly, this perturbation changed dentate excitability and impaired contextual discrimination only during the acute but not chronic phase. This apparent recovery could be due to delayed axon sprouting of interneurons shown in the chronic phase(Jinde et al., 2012). Such slow compensatory mechanisms would not be present in acute and specific manipulation, as performed in our study. Although the emphasis and particularities of these studies differ, an overall picture emerges pointing to a clear functional separation of dorsal and ventral MCs.

Even though entorhinal cortex contributes to context memory formation(Kitamura et al., 2015), full medial entorhinal lesions do not alter contextual fear conditioning(Hales et al., 2014). Likewise, deleting NR1 subunit of NMDA receptors in dentate granule cells does not affect contextual fear conditioning using different contexts, though only very similar contexts can cause discrimination problems in the mutant mice(McHugh et al., 2007). In agreement with our results, entorhinal inputs to DG only fine tune pattern separation while acquiring contextual memory, and direct entorhinal input to CA3 underlies contextual memory retrieval(Leutgeb et al., 2007; McHugh et al., 2007). However, direct optogenetic inhibition of dorsal GCs block context memory acquisition(Bernier et al., 2017; Kheirbek et al., 2013), which indicates that dorsal GCs firing is indispensable for context memory formation. Consistent with this evidence, our data demonstrates that memory acquisition requires ventral MCs input to dorsal GCs. As the ventral MCs projection spreads across most of the dentate gyrus, we propose that ventral MCs activity conveying novelty signals functions as a gain control mechanism for dorsal GCs activity during acquisition. This hypothesis is further supported by the finding that GCs dendrites have voltage attenuation, which can strongly weaken entorhinal synaptic input(Krueppel et al., 2011; Schmidt-Hieber et al., 2007). Yet, ventral MCs axons form synapses much closer to the GCs soma than those originating from the entorhinal cortex, so they are ideally positioned to regulate action potential firing in dorsal GCs(Scharfman, 2016).

Contrary to the currently accepted view that ventral and dorsal hippocampal subdivisions are functionally independent, our results demonstrate that novelty information relayed through a ventro-dorsal hippocampal connection links these subdivisions - a crucial interaction for contextual memory formation.

## Acknowledgments

We thank Peter Jonas and Peter Somogyi for critically reading the manuscript, Satoshi Kida for helpful discussion, Taijia Makinen for providing the Prox1-creERT2 mouse line and Hiromu Yawo for the VAMP2-Venus construct. We also thank to Vivek Jayaraman, Ph.D., Rex A. Kerr, Ph.D., Douglas S. Kim, Ph.D., Loren L. Looger, Ph.D., Karel Svoboda, Ph.D. from the GENIE Project, Janelia Farm Research Campus, Howard Hughes Medical Institute, for the viral constructs used for GCaMP6s expression. We also thank to Jacqueline Montanaro, Vanessa Zheden, David Kleindienst for technical assistance and Robert Beatti for imaging assistance.

## Author contributions

F.F. conceived the project and performed the experiments. F.F., M.J. and R.S. designed the experiments. A.S. performed the EM and in-situ hybridization experiments and assisted on surgery. P.K. performed the slice electrophysiology experiments. K.K. provided the virus AAV_1_-CAG-Vamp2-Venus and AAVDj-FLEX-rev-ChR2-HA-2a-Hm4Di. M.J. analyzed the calcium imaging data. F.F. analyzed all the other data. F.F., M.J. and R.S. wrote the paper.

---

**Figure S1. Related to Figure 1.**
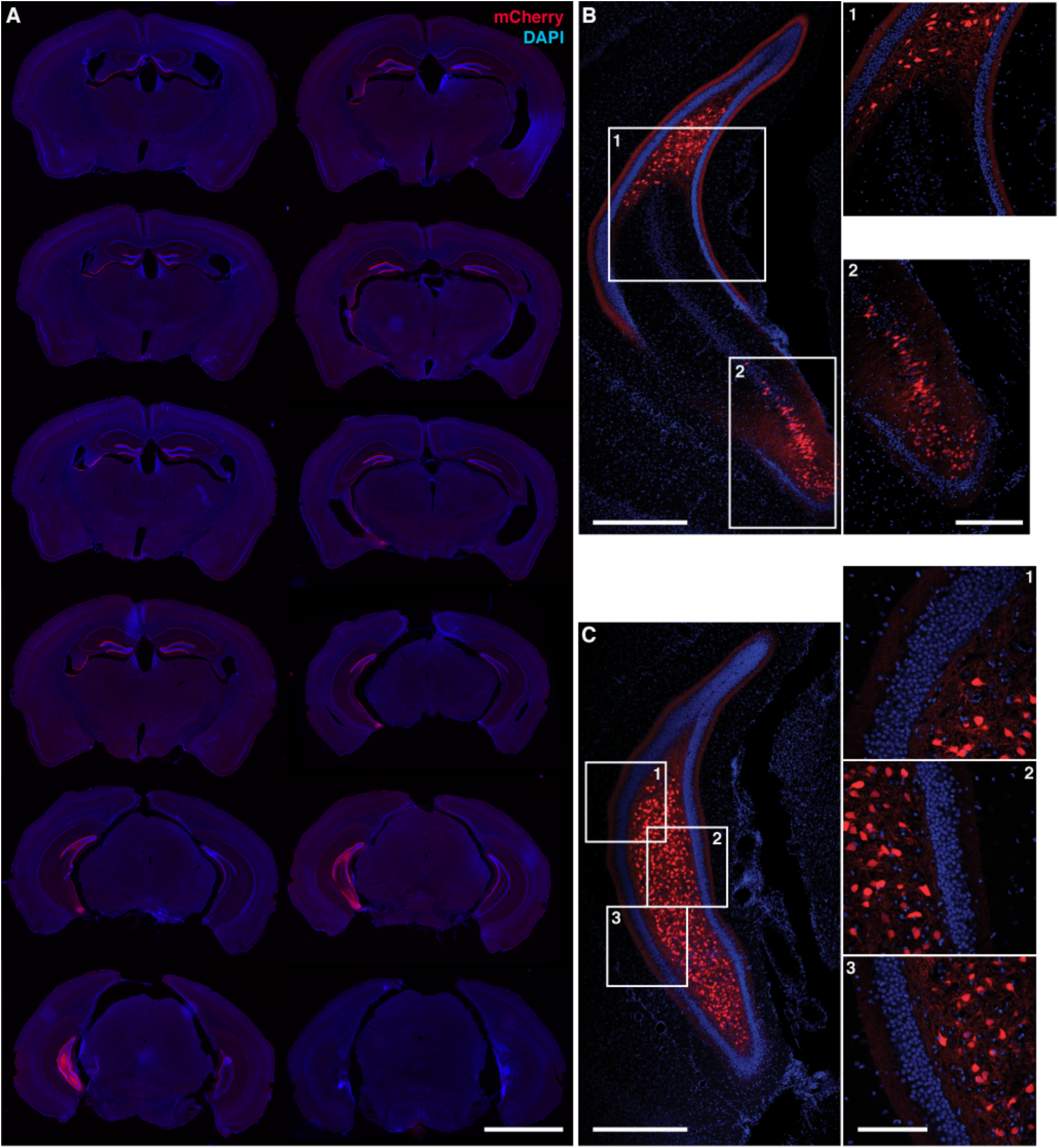
Extension of the ventral mossy cell axonal projections and distribution of mCherry-expressing cells in a Calb2-cre animal injected with AAV_5_-hSyn-DIO-mCherry into the ventral hilus. **A**. Rostro-caudal reconstruction of a unilateral injection of pAAV_5_-hSyn-DIO-mCherry into the ventral hilus of a Calb2-cre animal. Left upper corresponds to the most rostral part. Lower right is the most caudal. Scale bar: 2 mm. **B**. Section of intermediate dentate gyrus. Some mCherry-positive cells can be seen in the ventral CA3 area (Inset 2). Notice the complete absence of CA3 mCherry-postive cells in the dorsal intermediate portion (Inset 1) **C**. Ventral one third of the dentate gyrus. Only ventral hilar cells can be observed. Altogether, we found 417 ventral MCs and 33 CA3 cells (7,3 %) in all sections analyzed. No cells in the granule cell layer or molecular layer were found. Scale bars in **B** and **C** = 500 μm. Insets = 200 μm.

**Figure S2. Related to Figure 1.**
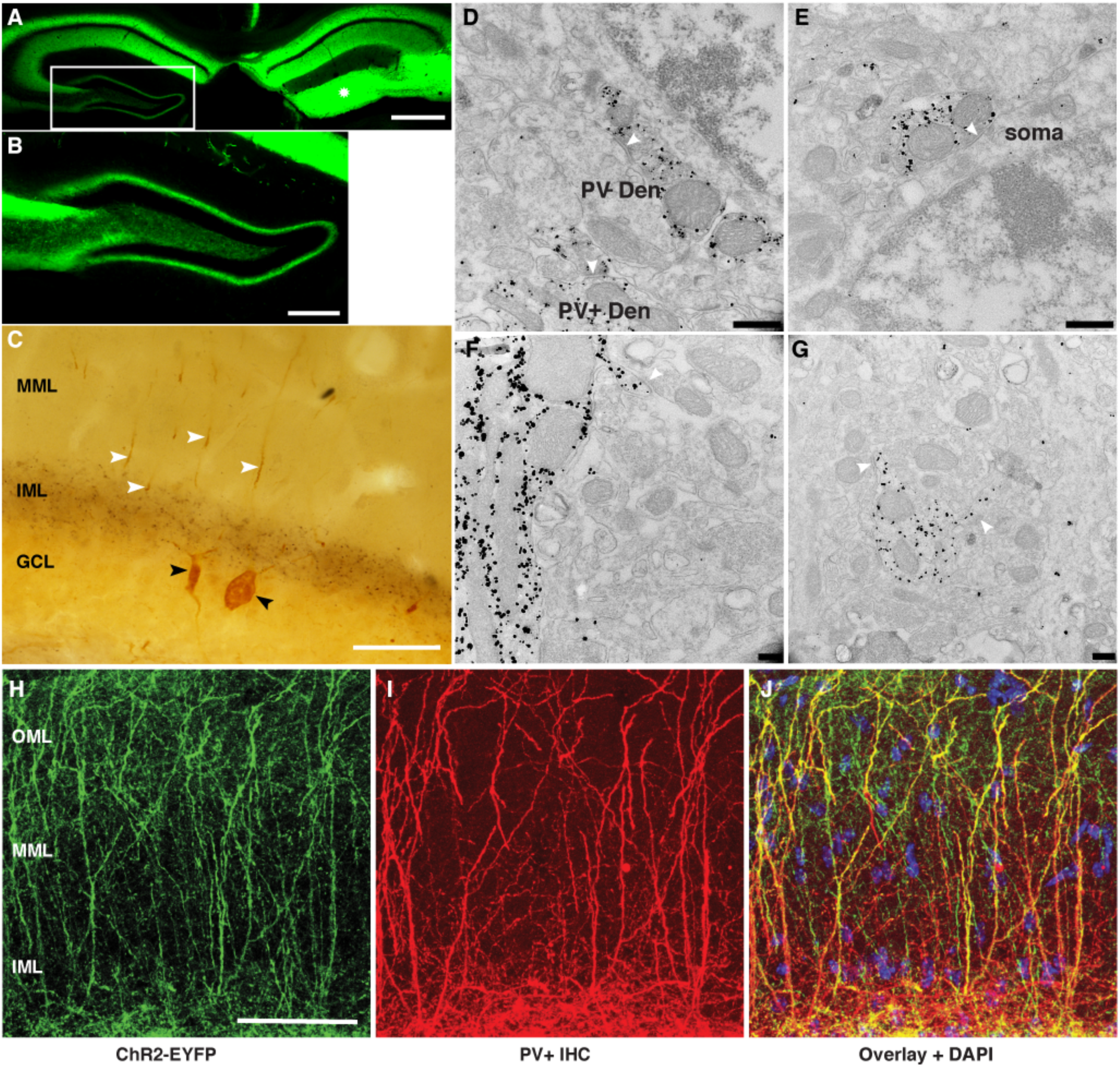
Double labeling of dorsal mossy cell axon terminals and PV-positive interneurons and quantification of PV+ dendrites in the molecular layer. **A.** Injection of pAAV_1_-CAG-VAMP2-venus into the dorsal dentate gyrus/CA3 (white asterisk) and its contralateral projection to CA3 and inner molecular layer of the dentate gyrus. Scale bar: 500 µm. **B.** Higher magnification of the area enclosed in the white rectangle in **A**. Scale bar: 200 µm. **C**. Photomicrograph of the dentate gyrus after pre-embedding labeling for VAMP2-venus (DAB) and PV (immunogold). Black arrowheads indicate the cell bodies of PV+ interneurons labeled with silver-intensified 1.4nm gold particles. White arrowheads indicate the ascending dendrites of these cells into the molecular layer. A dense band of DAB-positive boutons originating from the contralateral dorsal MCs can be seen in the inner molecular layer (IML). MML, medial molecular layer; GCL, granule cell layer. Scale bar: 50 µm. **D-E.** PV+ terminals labeled with silver-intensified 1.4 nm gold particles making symmetrical synapses (arrowheads) with PV+ dendrite (DEN), PV-dendrite (**D**) and soma (**D**). **F-G.** PV+ dendrites with spines making asymmetrical synapses (arrowheads) with non-labeled terminals. Scale bars: for **D** and **E** = 400 nm. For **F** and **G** = 1µm. **H-J**. Confocal images (maximum intensity projection) of ChR2-EYFP (**H**), immunoreactivity for PV (**I**) and overlay plus DAPI nuclear staining (**J**) in a vGAT-ChR2-EYFP mouse. Out of 47 EYFP+ dendrites, 40 were also positive for PV (85 %, n = 3 mice). OML, outer molecular layer; MML, medial molecular layer; IML, inner molecular layer. Scale bar = 100 μm.

**Figure S3. Related to Figure 2.**
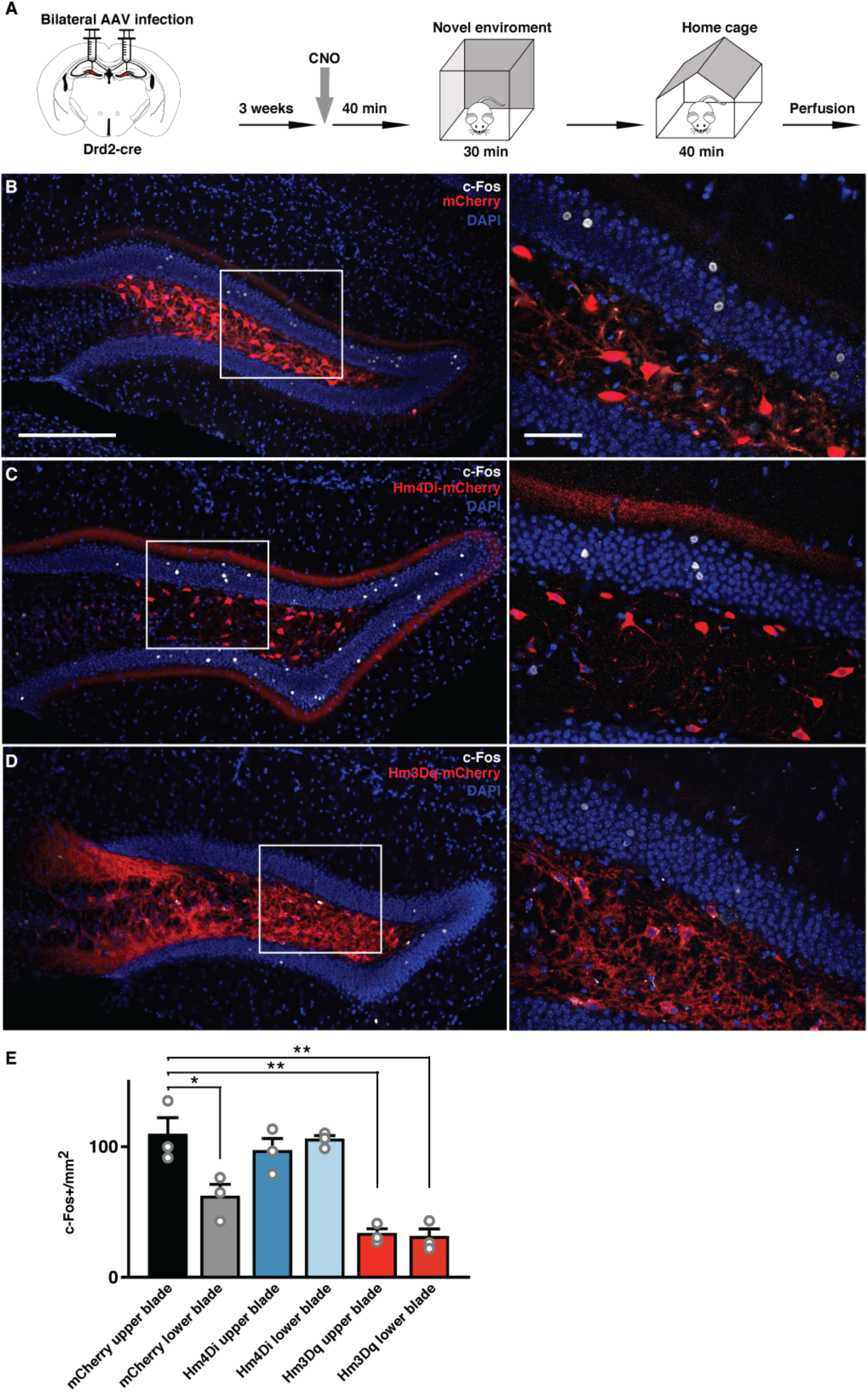
c-Fos expression in Drd2-cre mice after chemogenetic manipulation of dorsal MCs during novel environment exploration. **A**. Schematic of the behavioral paradigm. **B**. Representative image of the dorsal DG expressing mCherry in dorsal MCs (red). White rectangle indicates the area of the inset on the right. c-Fos-positive neurons labeled in white, same for **C** and **D**. **C**. Representative image of the dorsal DG expressing Hm4Di-mCherry in dorsal MCs (red). **D**. Representative image of the dorsal DG expressing Hm3Dq-mCherry in dorsal MCs (red). Note the decrease of c-Fos-positive cells in the granule cells layer compared with **B**. **E**. Quantification of the c-Fos-positive cells. (n = 3 animals per group; mean + s.e. ** p < 0.01, * p = 0.045, t-test). (Scale bar = 200 µm, Inset = 50 µm)

**Figure S4. Related to Figure 2.**
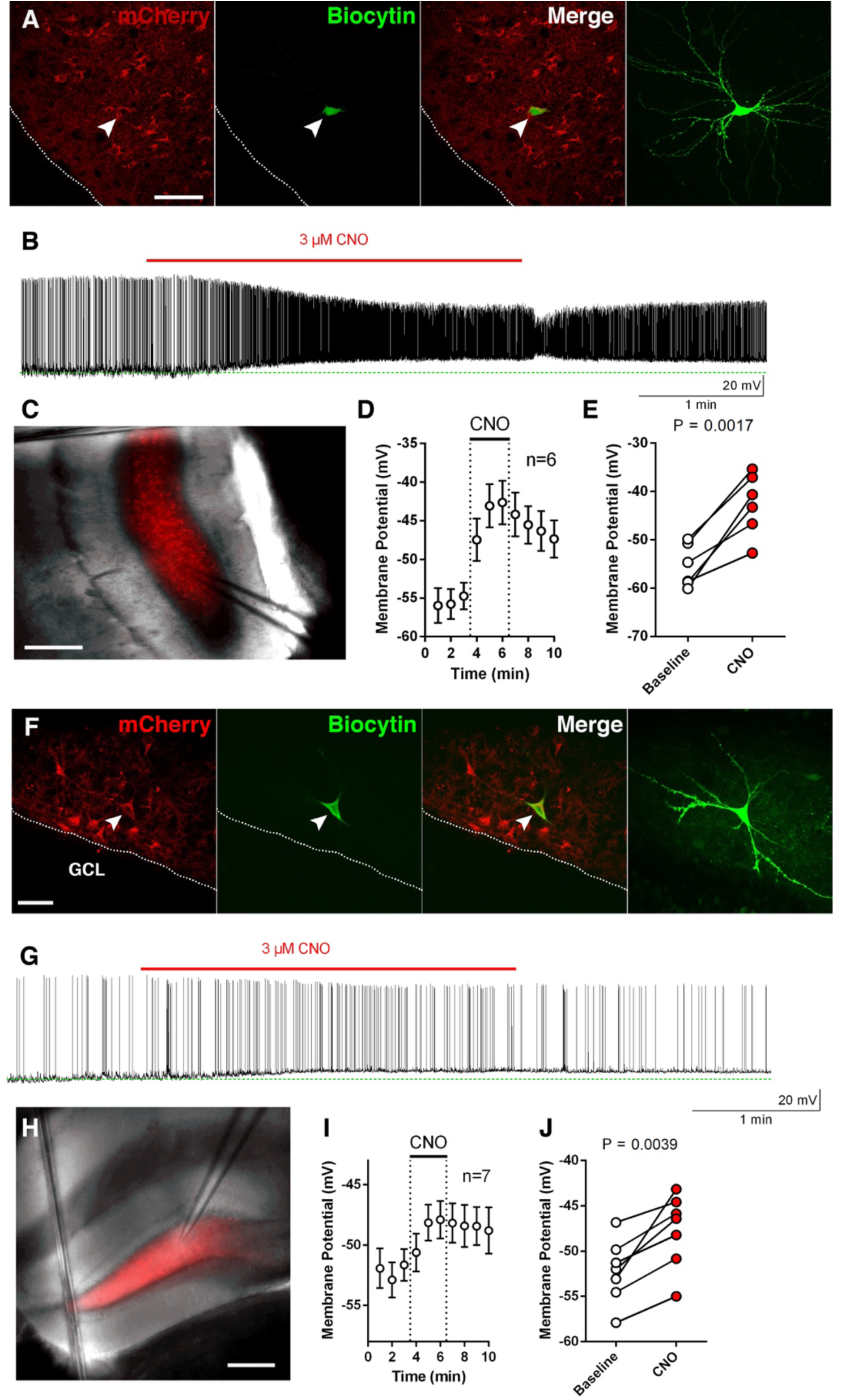
Hm3Dq-expressing ventral and dorsal MCs are depolarized by CNO. **A**. Example of an mCherry-positive, biocytin-filled ventral MC (green) in a Calb2-cre mouse injected with AAV5-hSyn-DIO-Hm3Dq-mCherry into the ventral hilus (scale bar = 50 µm). **B**. Example trace showing whole-cell current-clamp the depolarizing effect of CNO in an hM3Dq-expressing ventral MC. **C**. Example of the recording position of an mCherry-positive ventral MC. (scale bar = 200 µm) **D-E**. CNO significantly depolarized hM3Dq-expressing dorsal MCs (7 cells, 3 mice). **F**. Example of an mCherry-positive, biocytin-filled dorsal MC (green) in a Drd2-cre mouse injected with AAV5-hSyn-DIO-Hm3Dq-mCherry into the dorsal hilus (scale bar = 50 µm). **G**. Example trace of a whole-cell current-clamp recording showing the depolarizing effect of CNO in an hM3Dq-expressing dorsal MC. **H**. Example of the recording position of an mCherry-positive dorsal MC. (scale bar = 200 µm) **I-J**. CNO significantly depolarized hM3Dq-expressing dorsal MCs (7 cells, 3 mice). p-values were calculated with the paired t-test.

**Figure S5. Related to Figure 4.**
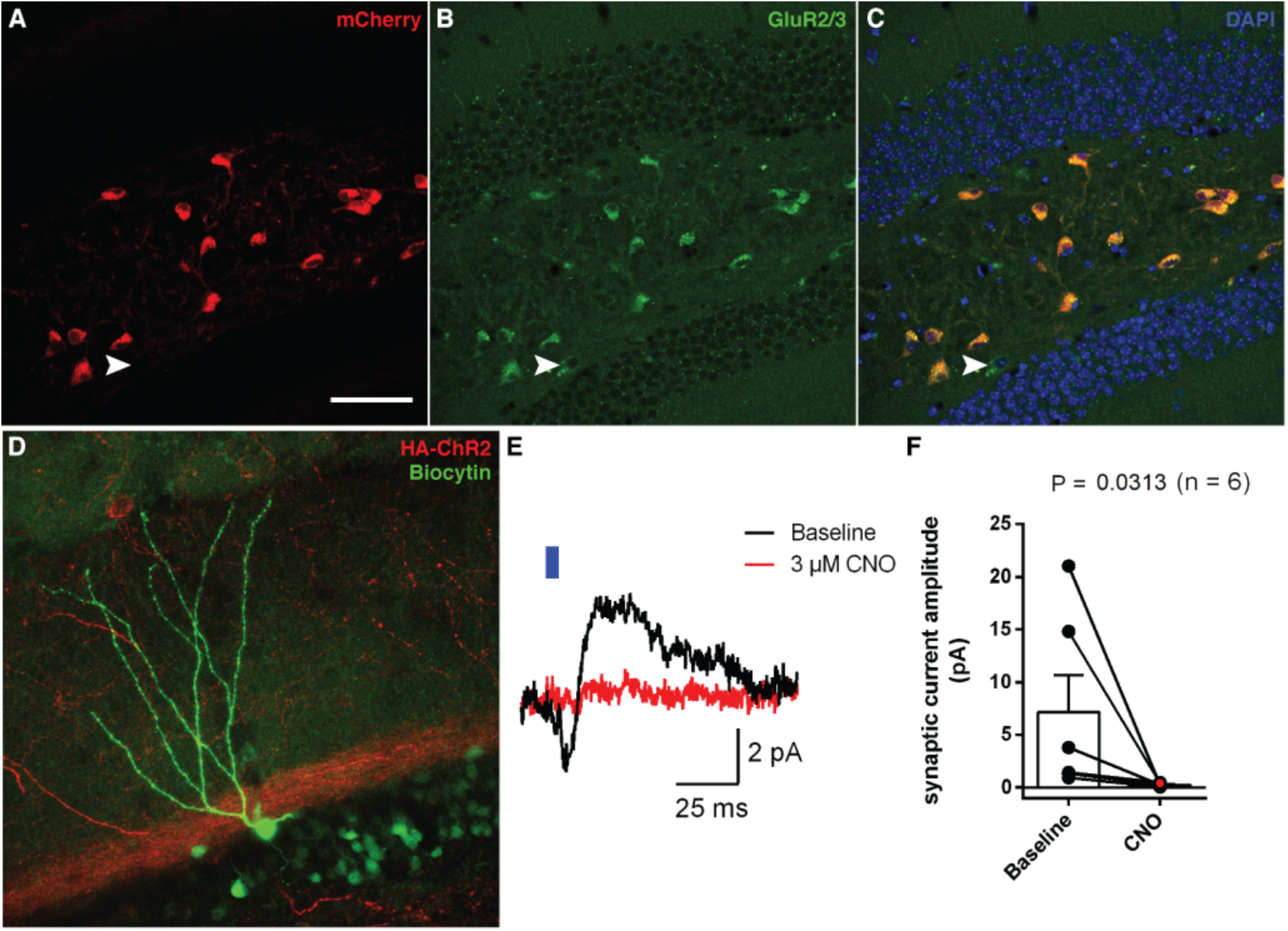
Quantification of dorsal MCs and suppression of light-evoked synaptic currents from dorsal MC to dorsal GCs via activation of pre-synaptic Hm4Di. **A.** Cells expressing mCherry (red) in dorsal hilus of a Drd2-cre mouse. **B**. Cells immunopositive for GluR2/3 (green). **C**. merge image plus DAPI (blue). Note that all mCherry positive cells are also GluR2/3 positive and only one GluR2/3 positive cell is negative for mCherry (white arrow head). Scale = 100 µm. **D**. Example of a biocytin-filled dorsal GC (green) and ChR2-positive projections originating from dorsal MCs (red) in a Drd2-cre mouse injected with AAVDj-FLEX-rev-ChR2-HA-2a-Hm4Di (scale bar = 20 µm). **E**. Example of a di-synaptic response evoked by a 5 ms light pulse before and after the application of CNO. Traces resulted from averaging 43 and 56 individual responses before and 5 min after the application of CNO, respectively. **F**. CNO significantly reduced the average maximal amplitude of light-evoked synaptic currents (6 cells, 2 mice). p-values was calculated with the Wilcoxon test.

**Figure S6. Related to Figure 6.**
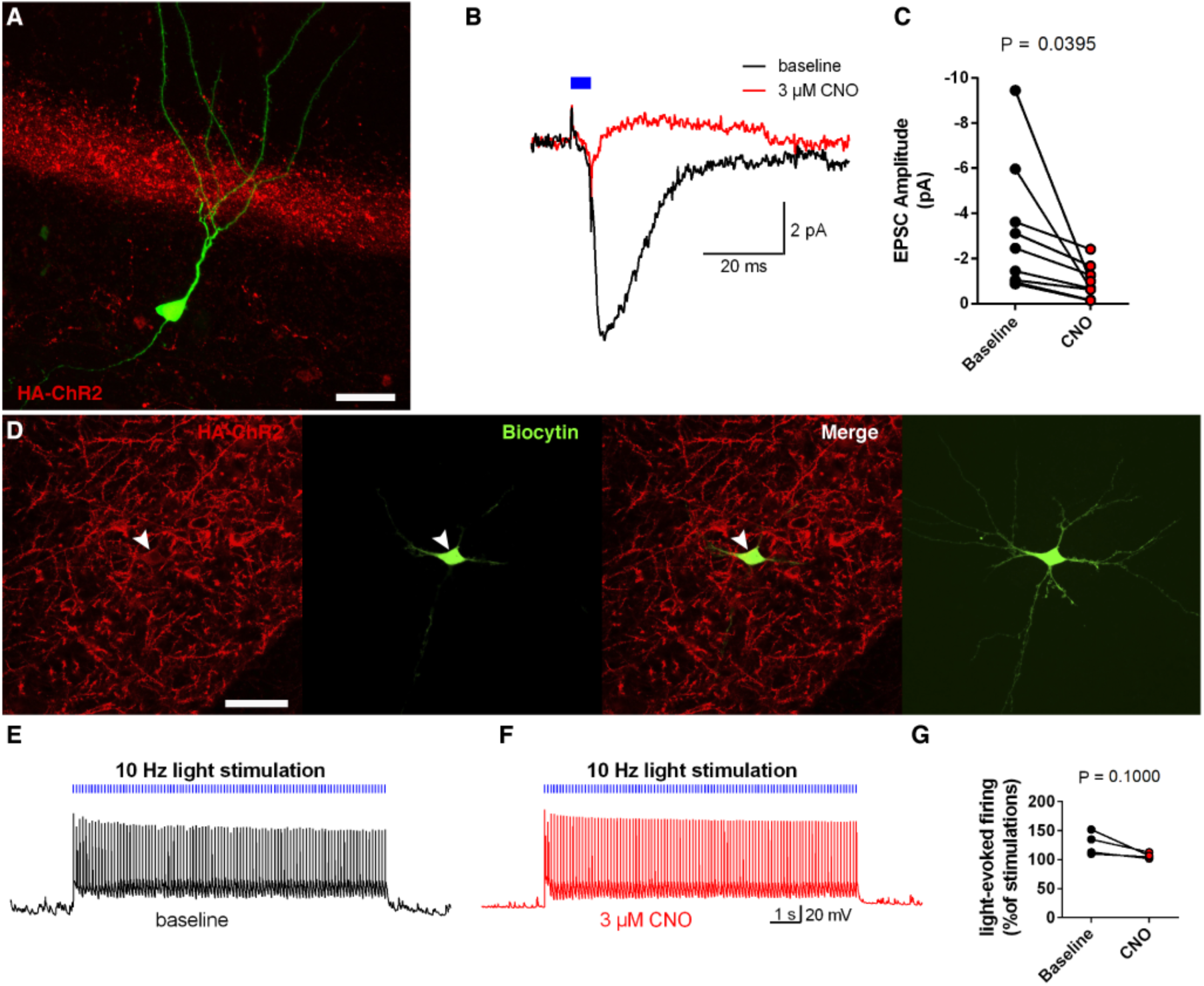
Suppression of light-evoked synaptic currents from ventral MC to dorsal GCs via activation of pre-synaptic Hm4Di. **A**. Example of a biocytin-filled dorsal GC (green) and ChR2-positive projections originating from ventral MCs (red) in a Calb2-cre mouse injected with AAVDj-FLEX-rev-ChR2-HA-2a-Hm4Di (scale bar = 20 µm). B. Example of EPSCs evoked by a 5 ms light pulse before and after the application of CNO. Traces resulted from averaging 52 and 51 individual responses before and 5 min after the application of CNO, respectively. C. CNO significantly reduced the average amplitude of light-evoked EPSCs (9 cells, 7 mice). D. Example of a ChR2-expressing (red), biocytin-filled ventral MC (green) (scale bar = 50 µm). E-F Example traces of light-evoked firing in a ventral MC before (E) and 5 min after the application of CNO (F). G. CNO application did not significantly affect the success rate of light-evoked action potential generation at the soma (4 cells, 2 mice). p-values were calculated with the paired t-test.

**Figure S7. Related to Figure 7.**
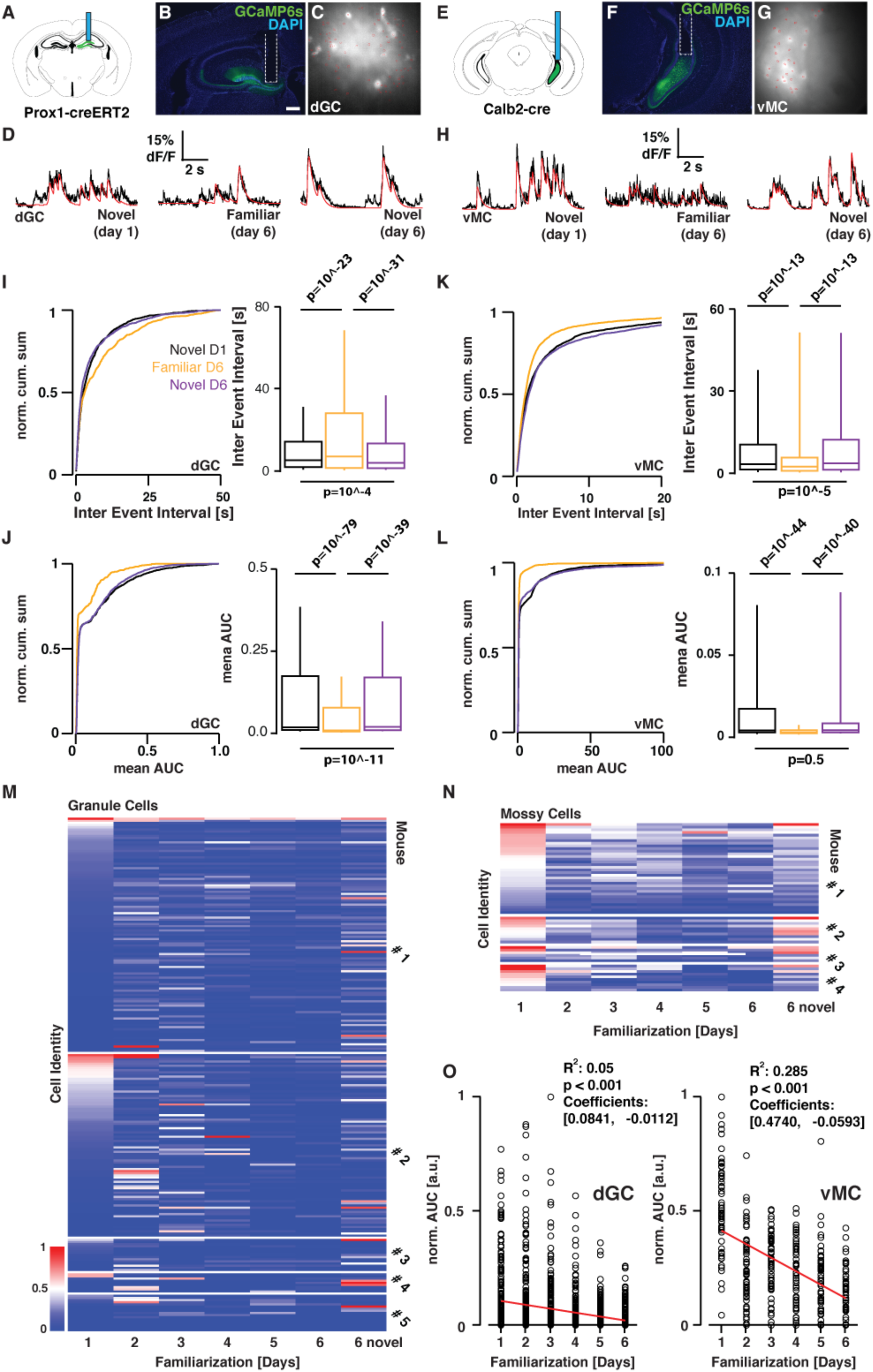
Compilation of Calcium imaging data. **A**. Prox1-creERT2 animals were unilaterally injected with AAV9-hSyn-Flex-GCaMP6s into the dorsal DG. Subsequently, a GRIN lens (blue rod) was implanted above the injection site. **B**. Example of a GRIN lens implantation site in a Prox1-creERT2 animal expressing GCaMP6s in granule cells in the dorsal dentate gyrus. **C**. Maximum projection image with superimposed manually selected ROI (red) of a 3 min long recording of dorsal GCs in one animal implanted as described in (**A**). **D**. Example dorsal GC Calcium responses recorded across familiarization and novel environments for the same animal (black traces). We used Markov Chain Monte Carlo method develop for spike inference in continuous time (see methods) and used this inference to generate fitted calcium traces (red traces). **E**. Calb2-cre animals were unilaterally injected with AAV5-hSyn-Flex-GCaMP6s into the ventral hilus. Subsequently, a GRIN lens (blue rod) was implanted above the injection site. **F**. Example of a GRIN lens implantation site in a Calb2-cre animal expressing GCaMP6s in ventral mossy cells in the ventral dentate gyrus. Higher magnification images of the framed areas in the upper images are shown in the bottom. Scale bars: 500 µm. **G**. as in (**C**) but for an animal implanted as described in (**E**). **H**. As in (**D**), but for ventral MCs responses. **I-L.** Normalized cumulative interevent intervals and AUC distributions for dorsal GCs (**I, J**) and ventral MCs (**K, L**). Inset: box-plot of the average AUC. Center line: median value; box limits: 25^th^ and 75^th^ percentile values; whiskers: 10^th^ and 90^th^ percentile. Black: 1^st^ day, yellow: 6^th^ day of familiarization; purple: 6^th^ day, but in novel environment. Statistics: p-values for the Kolmogorov–Smirnov test. Note that for ventral MCs, the decrease of average activity after familiarization is due to the change in event amplitude. For dorsal GCs, the change in population activity after familiarization is driven by the decrease in amplitude and frequency of the events (ventral MCs = 4 animals, 60 cells, dorsal GCs: 5 animals; 229 cells). **M-N**. AUC (normalized to the max. AUC per animal) for all recorded dorsal GCs (**M**) and ventral MCs (**N**) during the entire recording session (3 min). Each row depicts an ROI recorded over different sessions. All recorded cells are aligned in columns and are sorted by mean activity strength per ROI in the first recording session. Note: (i) To avoid biases due to changes in absolute ΔF/F we normalized each ROI to the maximal ΔF/F across all 7 recording sessions; (ii) As expected from dorsal GC place field properties, different populations of dorsal GCs get activated in different environments (compare column day 1 with column day 6 novel). In contrast, ventral MCs appear to be reactivated similarly in both environments (day1 as in day 6 novel), suggesting a role in novelty detection. **O**. Linear regression of the normalized AUC data points for the familiarization data point in **M** & **N** (normalized to maximal AUC for each group).

**Table S1. Related to Figure 1.** Raw data for EM synapse count. Areas examined in each animal were 1650-2520 µm^2^.

|  | DAB-positive synapses on PV-positive dendrites | DAB-positive synapses on GC spines | DAB-positive synapses on GC spines in total DAB-positive synapses | I<sub>p/g</sub> |
| --- | --- | --- | --- | --- |
| **Ventral MCs Animal 1** | 7.69% | 8.12% | 96.5% | 0.95 |
| **n=** | 26 | 727 |  |  |
| **Ventral MCs Animal 2** | 5.26% | 4.95% | 98.0% | 1.06 |
| **n=** | 19 | 909 |  |  |
| **Ventral MCs Animal 3** | 6.06% | 9.73% | 96.1% | 0.62 |
| **n=** | 33 | 822 |  |  |
| **Ventral MCs Animal 4** | 5.88% | 6.40% | 96.1% | 0.92 |
| **n=** | 34 | 829 |  |  |
| **Averages for Ventral MCs Animals (n=4)** | **6.22%** | **7.30%** | **96.7%** | **0.89** |
| **Dorsal MCs Animal 1** | 32.98% | 8.69% | 89.1% | 3.80 |
| **n=** | 94 | 771 |  |  |
| **Dorsal MCs Animal 2** | 24.53% | 8.09% | 93.4% | 3.03 |
| **n=** | 53 | 754 |  |  |
| **Dorsal MCs Animal 3** | 20.27% | 11.28% | 88.9% | 1.80 |
| **n=** | 74 | 594 |  |  |
| **Dorsal MCs Animal 4** | 16.67% | 7.00% | 96.2% | 2.38 |
| **n=** | 24 | 615 |  |  |
| **Averages for Dorsal MCs Animals (n=4)** | **23.61%** | **8.77%** | **91.9%** | **2.75** |

Movie S1. Related to Figure 3.

Locomotion and calcium imaging of dorsal granule cells of a freely behaving mouse 3 min and 30 min after CNO injection. Ventral mossy cells expressing Hm3Dq bilaterally.

Movie S2. Related to Figure 7.

Locomotion and calcium imaging of dorsal granule cells of a freely behaving mouse in novel environment on day 1, familiar environment on day 6 and novel environment on day 6.

Movie S3. Related to Figure 7.

Locomotion and calcium imaging of ventral mossy cells of a freely behaving mouse in novel environment on day 1, familiar environment on day 6 and novel environment on day 6.

## Contact for Reagent and Resource Sharing

Further information and request for resources and reagents should be directed to and will be fulfilled by the Lead Contact, Ryuichi Shigemoto.

## Data and Software Availability

All data and analysis code are available upon request from the Lead Contact Ryuichi Shigemoto.

## Experimental Models and Subject Details

Mouse protocols were reviewed by the institutional preclinical core facility (PCF) at IST Austria. All breeding and experimentation were performed under a license approved by the Austrian Federal Ministry of Science and Research in accordance with the Austrian and EU animal laws. Male mice C56BL6/J, Calb2-cre (JAX 010774), Drd2-cre (B6.FVB (Cg)-Tg (Drd2-cre)ER44Gsat/Mmcd, MMRRC), vGAT-ChR2-EYFP (JAX 014548) and Prox1-creERT2, developed by Taijia Makinen(Bazigou et al., 2011) were all on a C56BL6/J background and 2 to 8 months old for use in this study. Littermates were housed together before surgery. Animals received food and water *ad libitum* and were housed under a 12 h light-dark cycle. Littermates were randomly allocated to experimental groups. After surgery animals were housed individually and allowed to recover for at least 3 weeks before experiments.

## Materials and Methods

### Virus vectors

AAV5-hSyn-DIO-mCherry, AAV5-hSynDIO-Hm4Di-mCherry and AAV5-hSyn-DIO-Hm3Dq-mCherry vectors were obtained from University of North Carolina vector core and Addgene viral service. AAV9-hSyn-Flex-GCaMP6s-WPRE-SV40 and AAV5-hSyn-Flex-GCaMP6s-WPRE-SV40 were obtained from University of Pennsylvania Vector Core. pAAV1-CAG-Vamp2-venus transfer plasmid was generated by subcloning VAMP2-venus gene cassette obtained from Hiromu Yawo into pAAV-CaMKII-GFP (Addgene). Plasmid AAV-CAG::FLEX-rev::ChR2-HA-2a-hM4D was obtained from Addgene. Viral titrations were between 8 × 10^12^ and 7.5 × 10^13^ genome copy per ml. For inducing expression in Prox1-creERT2 animals, tamoxifen (T5648, Sigma) previously diluted in corn oil was injected i.p. (75 mg/kg body weight) for 5 consecutive days.

### Stereotactic injections, cannula and GRIN lens implantation

Animals were anaesthetized using a mixuture of 100 mg kg^−1^ Ketamine / 10 mg kg^−1^ Xylazine and injected 200 mg kg^−1^ of Novalgin as an intraoperative analgesic. Animals were placed in a stereotaxic apparatus (Kopf). The skull was exposed and craniotomies were performed at the following coordinates: ventral hilus injections or GRIN lens/cannula implantation; anteroposterior (AP) −3.2 mm, ±3.0 mm mediolateral (ML), dorsal dentate gyrus injections or GRIN lens/cannula implantation; −1.8 mm AP and ±1.3 mm ML. Virus injections were performed at a controlled rate of 0.1 µl min^−1^ using a water filled glass micropipette coupled to a 1 µl Hamilton microsyringe (7001; Hamilton). The pipette was slowly lowered to the target site and remained in place for another 5 min post-injection before being slowly withdrawn. For ventral hilus injections, 0.5 µl of undiluted virus was delivered at two dorso-ventral (DV) coordinates measured from the dura DV = −3.5 mm and DV = −3.7 mm. For dorsal dentate gyrus 0.7 µl of AAV_9_-hSyn-Flex-GCaMP6s-WPRE-SV40 diluted 1:7 in PBS was injected at DV = −1.9 mm below the dura. For mice used in intracerebral CNO infusions, bilateral guide cannulas were implanted above the dorsal dentate gyrus (−1.7 mm AP; ±1.0 mm ML; –1.7 mm DV) or two unilateral cannulas (PlasticsOne) were implanted above the ventral hilus on each hemisphere (−3.2 mm AP; ± 3.1 mm ML; −3.2 mm DV). For GRIN lens (DORIC lenses, Quebec) implantation four M1 screws were secured into the skull at the anterior and posterior edges of the surgical site to anchor the implant. After drying the surface of the skull, the lens was lowered slowly into the target, while imaging until the desired fluorescence was achieved. All cannulas and GRIN lens were secured to the skull using Super-bond dental cement (Sun medical, Japan). After surgery animals were injected with 5 mg kg^−1^ of Metacam (meloxicam) subcutaneously as a postoperative anti-inflammatory and painkiller. The animals recovered in a heat pad and then transferred to their home-cages and monitored closely for any discomfort signals for the three following days.

### Calcium imaging in behaving mice and data analysis

Recording sessions began 4 to 6 weeks after lens implantation. First, animals were adapted to the manipulation procedure and to carrying the microscope on their heads in their home-cages for 4 days. Once adapted they explored a square environment (35 cm by 35 cm) constructed of 40 cm high matt black painted walls, visual cues in one (colored tape) and removable floor (laboratory surface paper) for 20 min every day for 6 days. Three movies of 3 min each were taken at 0, 5 and 10 min after the beginning of the exploration. The paper floor remained in the environment during all familiarization sessions for each animal to keep their individual odor constant. On day 6, after exploring the familiar environment, animals were transferred to a different environment: a square box (20 cm by 20 cm) with two white and two vertical black and white striped walls. The floor had stainless steel rods of 5mm diameter, separated by 1 cm. Animals explored this environment for 10 min. Two movies of 3 min each were recorded during this time. Environments were carefully cleaned with 70 % ethanol between animals. For familiarization in the fear conditioning environment, animals explored the conditioning cage (17 × 17 cm, Ugo Basile SLR, Italy) for 10 min for 6 consecutive days. The cage was not cleaned in order to keep their individual odor constant. After the 10 min exploration during day 6, a single foot-shock 1sec, 0,65 mA was delivered. Immediately after, animals explored a novel environment (circular environment 20 cm diameter, 40 cm walls). For the running wheel experiments, animals were housed with a running wheel after implantation. For familiarization, animals visited a square environment (35 cm by 35 cm) constructed of 40 cm high matt black painted walls, visual cues in one (colored tape) and removable floor (laboratory surface paper) including their running wheel for 20 min every day for 6 days. On day 6, after exploring the familiar environment, animals were transferred to a different environment: a circular environment (40 cm diameter) constructed of 40 cm high matt black painted wall, visual cues (colored tape) and white plastic floor, together with their running wheel. Calcium-imaging data was acquired at 20 Hz, constant excitation power and exposure to compare fluorescent changes across sessions and corrected for motion artifacts using DORIC Neuroscience Studio Software. The rare artifacts that could not be corrected were excluded from the analysis. Imaged field of views were highly constant across days. Region of interest from single cells were carefully drawn by hand based on videos taken on different recording days. To denoise and estimate the most likely neuronal activity underlying the imaged traces, we used Markov Chain Monte Carlo method develop for spike inference in continuous time (MCMC-method) provided in the CaImAn toolbox (Giovannucci et al.). We subsequently used this inference to generate the estimated calcium traces. This approach is more precise than other denoising approaches and inherently defines uncertainty in all inferred transients. To compare the relative change across days, we normalized all inferred events for every drawn ROI across all days to the single maximum inferred ΔF (t)/F_0_ measured for each ROI. We then calculated the area under the curve (AUC) (as a measure of neuronal activity for the first three minutes during exploration of either familiar or novel environment. We took advantage of the inherent uncertainty estimation of the MCMC-analysis to determine which events are significant. AUCs were calculated for all sessions and ROIs. We identified a total of 229 independently active dorsal granule cell ROIs and 60 ventral mossy cell ROIs for the familiarization experiment. For visualization purposes, we normalized all AUCs to the maximal AUC measured of all cells (Figure 7 B&G). To compare the global activity change across animals, we compared the average AUC for all cells per animal (Figure 7 C&H). All traces were normalized to the maximal average AUC measured across days. Inter-event intervals and peak distribution were extracted from the MCMC inferred traces. Peaks were required to have at least 1 % (from normalized maximum for each ROI). The same analysis was performed for the experiments in which dorsal GC were imaged during chemogenetic ventral MC activation (n = 386 dorsal GC ROIs). Control (mCherry) and experimental (Hm3Dq) animals were continuously recorded after CNO injection. A strong wave of neuronal activity (Figure 3 B-C, Supplementary Movie S1) was recorded only in experimental animals 30-33 min after injection. To extract the locomotion from the behavioral movies, a custom written MATLAB (Mathworks) script was used to contrast enhance, background subtract and fit an ellipsoid on the tracked mouse. The center of mass of the ellipsoid was used as the tracking point (see Supplementary Movies – red square on mice). We separately calculated the distribution of locomotion speeds for novel (data from day 1 and day 6 novel) and familiar (data from day 5 and day 6) environments in ventral MC and dorsal GC recorded animals, respectively (Figure 7 D, I). To correlate locomotion and neuronal activity, we used all AUC events, including epochs of no activity, calculated for each ROI and binned them for different locomotion speeds. All novel (day 1 and day 6 novel) and familiar (day 5 and day 6) events were used. For the running wheel analysis, only AUC during constant locomotion on the running wheel were used (seven epochs with a total duration of 153 s and 400 s for novel and familiar environments, respectively). As we observed an increase in average speed in the first 3-6 mins of exploration in Hm3Dq expressing animals after CNO injection (Figure 3D), we hypothesized that this increase is caused by a distress response to the injection. In order to show this, three WT control mice were injected with 3 mg/kg CNO i.p. and immediately after transferred to the same environment Hm3Dq expressing animals explored.

### Fear conditioning and novel environment exploration

Animals were acclimated to handling for three days before behavioral assays. A fear conditioning system (Ugo Basile SLR, Italy), coupled to Ethovision software (Noldus, Wageningen) was used for contextual fear conditioning and analysis. During pre-exposure, animals explored the conditioning chamber for 3 min. 24 hrs later, animals were reintroduced to the conditioning cage explored for 3 min. Then, three consecutive 1 s foot shocks of 0.65 mA with a 1 min interval between them were delivered. 24 hrs later, animals were placed again in the conditioning chamber for a 3 min exploration period. 24 hrs later, animals explored a distinct novel environment B, which consisted of an opaque black painted plastic tube of 35 cm diameter and 40 cm high. This environment was located in a different soundproof box than the conditioning chamber. Before every context exploration, the conditioning chamber was odorized with 1 % acetic acid, and context B was odorized with 0.25 % benzaldehyde. Both environments were carefully cleaned with ethanol 70 % after every exploration. For conditioning in a familiar environment, animals explored the conditioning cage (17 × 17 cm, Ugo Basile SLR, Italy) for 10 min for 6 consecutive days. In order to keep animal’s individual odor constant, a piece of laboratory paper was kept under the grid floor during all the familiarization and conditioning procedure. During day 6, animals receive 0.75 mg/kg of CNO i.p. and returned to their home cages for 40 min. Then, animals explored the conditioning box for 10 min. After this period, three consecutive 1 s foot shocks of 0.65 mA with a 1 min interval between them were delivered. 24 hrs later, animals explored the conditioning box for 3 min and 24 hrs later, they explored a novel environment (circular environment 20 cm diameter, 40 cm high) for 3 min. Freezing was measured with activity tracking in Ethovision, with threshold of 500 ms for immobility detection. For novel environment exploration and following perfusion for c-Fos immunolabelling, animals were handled for 3 days before experiments to acclimatize to the experimenter and procedures. After CNO injection, animals were kept in their home-cages for 40 min before being transferred to a square box of 20 cm by 20 cm, with two white walls, and two walls with vertical black and white lines with white plastic floor. Inside the cage there was a running wheel and a styrofoam box 5 cm by 7 cm and 3 cm high. Animals explored this environment for 30 min, then returned to their home-cage and remained there for another 40 min before perfusion.

### CNO delivery

Clozapine-N-oxide (CNO, Tocris, Catalog number 4936) was dissolved in dimethylsulfoxide (DMSO) at 1 % final concentration, and then diluted with PBS to 1 mg/ml for intraperitoneal (i.p.) injections and with ACSF at 3 µM for intracerebral infusions. For i.p. injections, we delivered 3 mg/kg or 0,75 mg/kg of CNO, followed by behavior 40 min after the injection. For intracranial delivery, CNO was infused bilaterally into the dorsal or ventral dentate gyrus with 0.3 µl per hemisphere at a concentration of 3 µM using a 26-gauge stainless steel internal cannula (PlasticsOne). The internal cannula was connected with a 1 µl Hamilton syringe (7001; Hamilton) to control the injection rate at 100 nl min^−1^. The injection cannula was left connected for 5 min before removal to allow for diffusion. Finally, all behavior was performed 15 min following the drug infusion.

### Imunohistochemistry and *in situ* hybridization

For immunohistochemistry, mice were overdosed with 750–1000 mg kg^−1^ mixture of Ketamine, Xylazine and perfused transcardially with PBS, followed by 4 % cold paraformaldehyde (PFA) in PB. Extracted brains were kept in 30% sucrose in PB at 4 °C overnight, then transferred to PBS. A sliding microtome (Leica SM2000R) was used to cut 40-µm coronal slices in PBS. Slices were washed with PBS-T (PBS + 0.2 % Triton X-100), then incubated with PBS-T + 20 % normal donkey serum, at room temperature for 1 h for blocking. Then, slices were incubated with one or more primary antibodies at 4 °C for 24 hr (rabbit anti-calretinin AB5054, Millipore, dilution 1:1000; goat anti c-Fos SC-52, Santa Cruz Biotechnology, dilution 1:500; rat anti-somatostatin Mab 354, Millipore, dilution 1:50, rabbit anti-Glutamate receptor GluR2/3, Chemicon AB1506, dilution 1:500). Three washes of PBS-T for 10 min each were performed on the slices before 1 h incubation with secondary antibody at 1:1000 dilution (A-21206; A-21447; A-21208 ThermoFisher Scientific). Slices were washed three more times in PBS-T for 10 min each, stained with 4ʹ,6-diamidino-2-phenylindole (DAPI) to label cell nuclei and mounted with Prolong antifade reagent (Thermofisher) onto microscope slides. For *in situ* hybridization, calb2-cre animals were injected hilus with pAAV5-hSynDIO-mCherry in the ventral hilus. Three weeks later, they were anesthetized with isofluorane, decapitated and their brains removed immediately. The ventral hippocampus was dissected, placed in a Cryomold (Sakura, Japan) filled with O.C.T. Tissue-Tek compound (Sakura, Japan), and stored at −80 °C until 14 µm sections were cut with a cryostat (Microm HM 560, Thermo) and mounted in Superfrost plus slides (Thermo scientific). The whole *in situ* hybridization procedure was performed with and according to the protocol of the RNAscope Fluorescent Multiplex Assay (ACD) using probes for vGluT1 and mCherry. Slides were imaged in an upright Zeiss LSM 700 confocal microscope.

### c-Fos counting

For counting c-Fos-positive granule cells in the dentate gyrus, three 40 µm sections were taken from dorsal DG and three for ventral DG from each animal if required. These sections were separated each other by 160 µm, thus representing the different antero-posterior levels of the whole dorsal or ventral DG. Pictures were taken in an upright Zeiss LSM 700 confocal microscope with the same magnification, laser power and exposure times. Then, the total area of the granule cell layer was measured, and cells positive for c-Fos within this area were counted using Fiji (https://fiji.sc). Finally, the density of cells was calculated dividing the number of cells by the area.

### Electron microscopy

Animal anesthetization, fixation and pre-embedding immunoelectron microscopy (EM) were performed as described previously(Parajuli et al., 2012). Briefly, mice were deeply anesthetized and trans-cardially perfused at a flow rate of 7 ml/min, first with 25 mM PBS for 1 min, followed by 10 min of fixative containing 4 % paraformaldehyde, 15 % picric acid and 0.05 % glutaraldehyde in 0.1 M PB. Brains were removed and kept in PB until coronal sections of 50 µm thickness were cut in a vibratome (Linear Slicer Pro7, D.S.K, Japan). Slices were washed in PB twice, cryo-protected, and freeze-thawed before adding blocking solution (10 % NGS in 2 % BSA) for 90 min. Double immunolabeling was done by mixing primary antibodies in 2 % BSA, incubated for 3 nights at 4 °C. For the ventral mossy cell samples, we mixed rabbit anti-PV (1:1000, Abcam) with mouse anti-RFP IgG1k (1:1000, MBL). For the dorsal mossy cell samples, we mixed rabbit anti-PV (1:1000, Abcam) with mouse anti-GFP IgG2a (1:3000, Wako). After washing, sections were incubated overnight at 4 °C in a mixture of secondary antibodies, 1.4 nm gold conjugated goat anti-rabbit (1:200, Nanoprobes) and goat anti-mouse IgG1k biotinylated (1:400, Life Technologies) antibodies for ventral mossy cell samples, and goat anti-mouse IgG2a biotinylated (1:10000, Life Technologies) antibodies for dorsal mossy cell samples. After 10 min post-fixation in 1 % glutaraldehyde we performed silver enhancement of immunogold particles (HQ Silver enhancement kit, Nanoprobes) and immunoperoxidase reaction, followed by 15 min fixation with 0.5 % osmium tetroxide and counterstaining with 1 % uranylacetate for 30 min. Slices were dehydrated in an ascending ethanol series, propylene oxide and then infiltrated in resin (Durcupan ACM, Sigma) at room temperature overnight. Slides were flat-embedded and resin was polymerized for 48 hrs at 60 °C. We have chosen the areas to be examined by the presence of the PV+ dendrites that were located in the inner one third of the molecular layer in a middle part of the upper blade of the dorsal dentate gyrus. The selected regions were re-embedded, and ultrathin serial sections at 70 nm thickness were cut with an ultramicrotome (Leica EM UC7) for examination in a transmission electron microscope (Tecnai 10, Philips, The Netherlands). The vast majority of DAB-positive synapses were made on GCs spines (Table S1, > 97% for ventral MCs, > 92% for dorsal MCs: Note that these values are minimum estimates because we targeted PV+ dendrites, making positive biases for synapses made on these dendrites over those on GCs) as described previously(Blasco-Ibáñez and Freund, 1997). We used a total number of 450 sections covering total examined area of 16,700 μm^2^. To estimate the relative contribution of ventral and dorsal MCs to synaptic innervation onto PV+ dendrites and GCs spines, we first calculated % of DAB-positive synapses in all synapses made on PV+ dendrites examined, and % of DAB-positive synapses in all synapses made on GCs spines in surrounding areas with similar depth from the section surface (Table S1). Then, these % values on PV+ dendrites were normalized by the % values on GCs spines in each animal to calculate an index Ip/g (Table S1), which indicates innervation strength on PV+ dendrites relative to that on GCs by MCs. The Ip/g values are independent of the fraction of transfected MCs. We compared Ip/g values between ventral and dorsal MCs samples with Student’s t-test (n = 4 animals for each).

### Slice electrophysiology

Drd2-cre or Calb2-cre mice (2.5 to 14 months old) were injected bilaterally with AAV expressing Hm4Di and ChR2 or Hm3Dq-mCherry in a Cre dependent manner. Three weeks post-injection they were anesthetized via i.p. injection of ketamine/xylazine (ketamine: 100 mg/kg; xylazine 10 mg/kg) and transcardially perfused with oxygenated (95 % O_2_ / 5 % CO_2_) ice-cold artificial cerebrospinal fluid (ACSF) containing (in mM): 118 NaCl, 2.5 KCl, MgSO_4_ 1.5, NaH_2_PO_4_ 1.25, CaCl_2_ 2, Myo-inositol 3, Glucose 10, Sucrose 40, NaHCO_3_ 25, pH = 7.4. The brain was rapidly excised and placed in a cutting chamber. Coronal brain slices were prepared with a D.S.K. Linear Slicer Pro7 (Dosaka, Japan) at 250 – 350 µm thickness for recordings of dorsal and ventral MCs. For recordings of optogenetically evoked currents in dorsal GCs, parasagittal slices were prepared at a 15-20 °C angle and at 1 – 1.5 mm thickness(Vega-Zuniga et al., 2018). Slices were left to recover for 20 min at 35 °C, thereafter heating was turned off and slices recovered to room temperature over the course of one hour. One slice was transferred to the recording chamber and superfused with recording ACSF (2 mM CaCl2) at a rate of 2-4 ml/min.

Whole-cell patch clamp recordings were performed using an EPC 10 USB amplifier with PatchMaster software (HEKA) at a sampling rate of 20 – 50 kHz and filtered at 2.9 kHz. MCs or dentate gyrus GCs were identified morphologically. Patch pipettes were pulled with a Model P-97 horizontal puller (Sutter Instrument, USA) at a resistance of 3 – 5 MΩ and filled with internal solution containing (in mM): K-Gluconate 140, MgCl2 2, HEPES 10, EGTA 0.5, MgATP 2, NaGTP 0.2, Sucrose 32, pH = 7.4 adjusted with KOH. In addition, 0.25 - 0.3 % biocytin was added to the internal solution for labelling of recorded neurons after slice fixation (4% paraformaldehyde). For optogenetic experiments, blue light (465 nm) was emitted from an LED (Plexon) and transmitted onto the slice through an optical fiber. Light pulses (5 ms duration, 15 mW) were applied at 10 Hz (for 10s at 30s interval) for current clamp experiments in MCs and at 0.1 - 0.5 Hz for voltage-clamp experiments in dorsal GCs. In voltage-clamp experiments, cells were held at −70 mV and light-evoked synaptic currents were measured by averaging traces of 50 - 100 light-stimulations before and 3 - 10 min after the start of the application of 3 µM CNO. To measure the effect of Hm3Dq activation on spontaneous action potential firing, the membrane potential was passively monitored. To test the effect of Hm4Di activation on light-evoked action potential firing, cells were hyperpolarized (∼-60-70 mV) to minimize spontaneous firing. Series resistance was monitored throughout all measurements and recordings with changes in series resistance exceeding 20% were discarded. Electrophysiological data was analyzed using Patchmaster software (HEKA), Clampfit 10 (Molecular Devices) and Prism 6 (GraphPad).

### Quantification and statistical Analysis

No statistical methods were used to predetermine sample size because means, variances and effect sizes were not predictable prior to performing the experiments. Thus, we decided to use five animals for behavioral experiments according to the number of animals commonly used for similar studies. After the experiments, we retrospectively performed the power analysis using means, variances and effect sizes obtained from our experiments, setting p < 0.05 and power of 0.9. For all the statistical comparisons that showed significant difference (p < 0.05), we confirmed that sample sizes were equal to or larger than those calculated by the power analysis. The investigators were not blind to group allocation.

